# Dispersal and Plant Arrangement Condition the Timing and Magnitude of Coffee Rust Infection

**DOI:** 10.1101/2022.04.07.487552

**Authors:** Emilio Mora Van Cauwelaert, Cecilia González González, Denis Boyer, Zachary Hajian Forooshani, John Vandermeer, Mariana Benítez

## Abstract

One central issue in coffee-leaf rust (*Hemileia vastatrix*) epidemiology is to understand what determines the intensity and the timing of yearly infections in coffee plantations. However, most experimental and theoretical studies report infection as an average at the plot level, obscuring the role of potentially key factors like rust dispersal or the planting pattern. Here, we first review the rust epidemic patterns of different sites, which reveal large variability in the duration and magnitude of the different epidemiologic phases. We then present a spatially explicit and parametrised model, where the host population is subdivided into discrete patches linked through spore dispersal, modelled as simple diffusion. With this model, we study the role of the planting arrangement, the dispersal intensity and plant-level variables on the maximum average tree infection (MATI) and its timing. Our results suggest that the epidemic timeline can be divided into two phases: a time lag and a growth phase *per se*. The model shows that the combination of the dispersal magnitude and plant aggregation modifies the MATI and the time to MATI, mainly by preventing some plants from reaching their maximum peak during the epidemic. It also affects the epidemic curves, which can have a stepped, or a rather smooth pattern in plots with otherwise similar conditions. The initial rust infection modulates the time lag before the epidemic and the infected leaf-fall rate drastically changes the MATI. These findings highlight the importance of explicitly considering the spatial aspects of coffee agroecosystems when measuring and managing rust infection, and help us to further understand the spatio-temporal dynamics of ecological systems in general.

## 1. Introduction

The first recorded epidemic of the coffee rust disease, caused by the fungus *Hemileia vastatrix*, broke out in Ceylon (now Sri Lanka) in 1869 (Talhinhas et al., 2017). Since then, coffee rust has spread across the continents, reaching virtually all the coffee plantations areas on earth (Avelino et al., 2006; McCook and Vandermeer, 2015). This is particularly relevant for farmers who depend economically on these crops.

One central issue in coffee rust epidemiology is to understand what determines the intensity of a one-year infection (Avelino et al., 2006; Gagliardi et al., 2020; Kushalappa and Eskes, 1989, Motisi et al., 2022). Similarly, many epidemiologists have sought to estimate the time to maximum infection for the design of specific control practices (Ananth, 1969; Burdekin, 1964). Both questions have been studied within the “disease triangle” framework (Stevens, 1960). In this sense, scientists and farmers have studied the pathogen’s properties such as its genetics (Carvalho et al., 2011), the host resistance and phenology (Avelino et al., 1993; Silva et al., 2006), and disease environmental drivers such as the temperature, humidity or precipitation (Avelino et al., 2015). Nevertheless, there is a large amount of variability in the intensity and timing of the different epidemiological phases of the coffee rust epidemic that remains unexplained, even between neighbouring coffee plots with the same environmental and biotic conditions (Li et al. 2022). For example, in some plantations, when the abiotic and biotic conditions for rust invasion are met, the infection may not start right away: there is a highly variable time lag (Boudrot et al., 2016; Mulinge and Griffiths, 1974).

To explain this variability, previous research has mainly focused on the infection phase *per se* (where susceptible leaves are infected by coffee rust spores and become infective; Talhinhas et al., 2017) and has overlooked another epidemiological phase: dispersal. Dispersal is the process by which spores are transported from one place to another, across different scales, ranging from the intraleaf, interleaf, or even the interplot scale (Becker and Kranz, 1977; Boudrot et al., 2016; Vandermeer and Rohani, 2014). Overall, the intensity of an epidemic is highly related to the rates of infection and dispersal during each season (Avelino et al., 2015). Analysis of dispersal is thus called for, and, by its very nature, must incorporate a spatial approach (Avelino et al., 2012). There are effectively two distinct scales of dispersal: first, the large scale mediated by the wind and resulting from a “rain” of spores over large areas (between plots or between farms; Becker and Kranz, 1977; Bowden, 1971; Kushalappa and Eskes, 1989), and second, the local scale, corresponding to neighbouring plants in a plot and/or leaves on the same plant, caused mainly by insect vectors, splash, wind gusts, and human action during harvest (Becker and Kranz, 1977; Vandermeer and Rohani, 2014). In this work, we will focus on the local plant-to-plant dispersal scale in a plot.

Intuitively higher local dispersal should lead to more severe epidemics in a plot. However, despite the importance of a nuanced understanding of rust epidemic dynamics within plots, most empirical studies on coffee rust epidemics report rust prevalence in terms of averages within a plot and not on individual trees (Bock, 1962a; Burdekin, 1964). This method reduces the sampling errors and smooths the epidemic curves but may obscure the relationship between plant and plot dynamics mediated by dispersal. Besides, coffee plantations can be arranged in rows or follow a more random arrangement depending on the age, type (rustic or conventional), or size of the plantation (Hajian-Forooshani and Vandermeer, 2021; Moguel and Toledo, 1999). Therefore coffee rust dispersal effect might be modulated by these planting patterns (Hajian-Forooshani and Vandermeer, 2021; Vandermeer et al., 2018). Finally, the relative importance of the dispersal between plants and the plant-level infection dynamics (such as the initial infection or the rust-infected leaf-fall rate) on the plot-level rust epidemics, has not been fully assessed or considered in current dynamic models (but see (Park et al., 2001)).

A dynamical modelling approach at both the plant and plot scales can help to disentangle such multi-scaled relations and processes, and shed light on the role of dispersal on the maximum infection prevalence (we will refer here to maximum infection) and the time to reach it in coffee plantations. We hypothesise that spatial dynamics and coffee rust dispersal might also play a role in the variability of the timing and magnitude of the different rust epidemiological phases in coffee plots with otherwise similar biotic and abiotic conditions. We thus seek to explore the determinants of a) the plot-averaged maximum infection and b) its timing, using a parametrised epidemiological SIX (Susceptible-Infected-eXternal inoculum) model in a spatially structured host population. With this model we analyse the role of the intensity of rust dispersal, planting arrangements, initial infection, and plant-level dynamics (such as the fall rate of an infected leaf) on the epidemiological outcomes.

## 2. Methods

We first reviewed quantitative and qualitative data on the maximum infection and the number of days to reach this maximum in different coffee systems. From these data, we estimated the duration of the different coffee rust epidemiological phases and their variability, and used them for future validation of the model. Secondly, we built a spatially explicit model to study the role of planting arrangement, dispersal intensity, plant-level dynamics, and initial conditions on the variability detected in the spatially-averaged maximum coffee rust infection and its timing. Details are presented below, but the overall work route was the following: We first constructed the model and parametrised it. Then, we implemented different computational scenarios at the plant and plot level, and studied their effect on the maximum infection, growth phase and time lags.

### 2.1 Reviewing the qualitative dynamics of coffee rust infection

To study the qualitative behaviour of coffee rust infections, we reviewed literature reporting data on coffee rust at coffee sites with different climatic, orographic, and management conditions. We used data from studies that presented at least a 12 month time series, rainfall pattern, and, in some cases, the harvesting period. We explored publications reporting all these data from the 1960’s to the present. Since most of the reviewed works do not report Tables or do not follow a uniform procedure, we extracted both the rainfall and coffee rust infection data directly from the graphs, using WebPlotDigitizer (Drevon et al., 2017; Rohatgi, 2020). We transformed all dates to their corresponding julian day number and all the precipitation histograms to millimetres. Some graphs started on day 250 (September 7th) and others on day 1 (January 1st) (Fig. 2). We considered three time series from Chiapas, México (Avelino et al., 1991; Vandermeer et al., 2018); two from Central America (Quetzaltenango, Guatemala and Turrialaba, Costa Rica) (Avelino et al., 1993; Boudrot et al., 2016); one from South India (Mysore) (Ananth, 1969); and three from Kenya (East Riff, Ruiru and Kiambu) (Becker and Kranz, 1977; Bock, 1962a; Mulinge and Griffiths, 1974). In the case of Ruiru, Kenya, rainfall was not directly reported, so we used the rainfall pattern reported for Nairobi, which has similar climatic conditions (Ndolo et al., 2017).

In some cases, coffee rust infection was reported as the average percentage of rust-infected leaves per tree, in others, as spores counted per tree or spores in the immediate vicinity. Rainfall was reported on a daily or monthly basis, and we grouped the daily data in months. When the harvesting period was reported, we added it manually, adding a ten day-error to each end. For each site, we reviewed the duration of the different epidemiological phases, the maximum infection reached and their relationship with rainfall and harvesting period. We defined the beginning of the rainy season as the middle of the first month that averaged precipitations higher than 50 mm, and the beginning of the epidemic growth period *per se* as the moment when the increase in the reported percentage of infected leaves in the trees was higher than 0.1 per day. The time between these two points was defined as the time lag. Other characteristics of the plantations such as the climatic, orographic, and management conditions are included in the supplementary material (S2). The general timings and magnitudes of infection on these sites were also compared to the predictions of our parametrised model.

### 2.2 SIX model construction

We modelled the dynamics of multiple individual plants, including both infection and internal leaf-to-leaf spore dispersal processes, coupled in a plot through a plant-to-plant dispersal mechanism. The overall modelling strategy is shown in Fig. 1 and can be summarised as follows. We first defined a 10 × 10 lattice (100 cells) with sink boundaries and 50 trees (N) in four different scenarios of planting arrangements: aggregated, random, rows, and spaced (Fig. 1B). These arrangements aim to represent the different patterns reported in coffee plots, where trees can be closely surrounded by other trees (aggregated), or have direct neighbours in one specific direction (rows), or have no direct neighbours (spaced) or have a random arrangement. Black squares represent the trees with susceptible or infected leaves as well as the immediate space around the tree where external uredospores may be present. White squares represent regions of space where only uredospores might be present (*S=I=0* in equation **2.2**) (Fig. 1B). We characterised the four different planting arrangements with a distance-between-plant index *<H>* defined as:

**Fig. 1.**
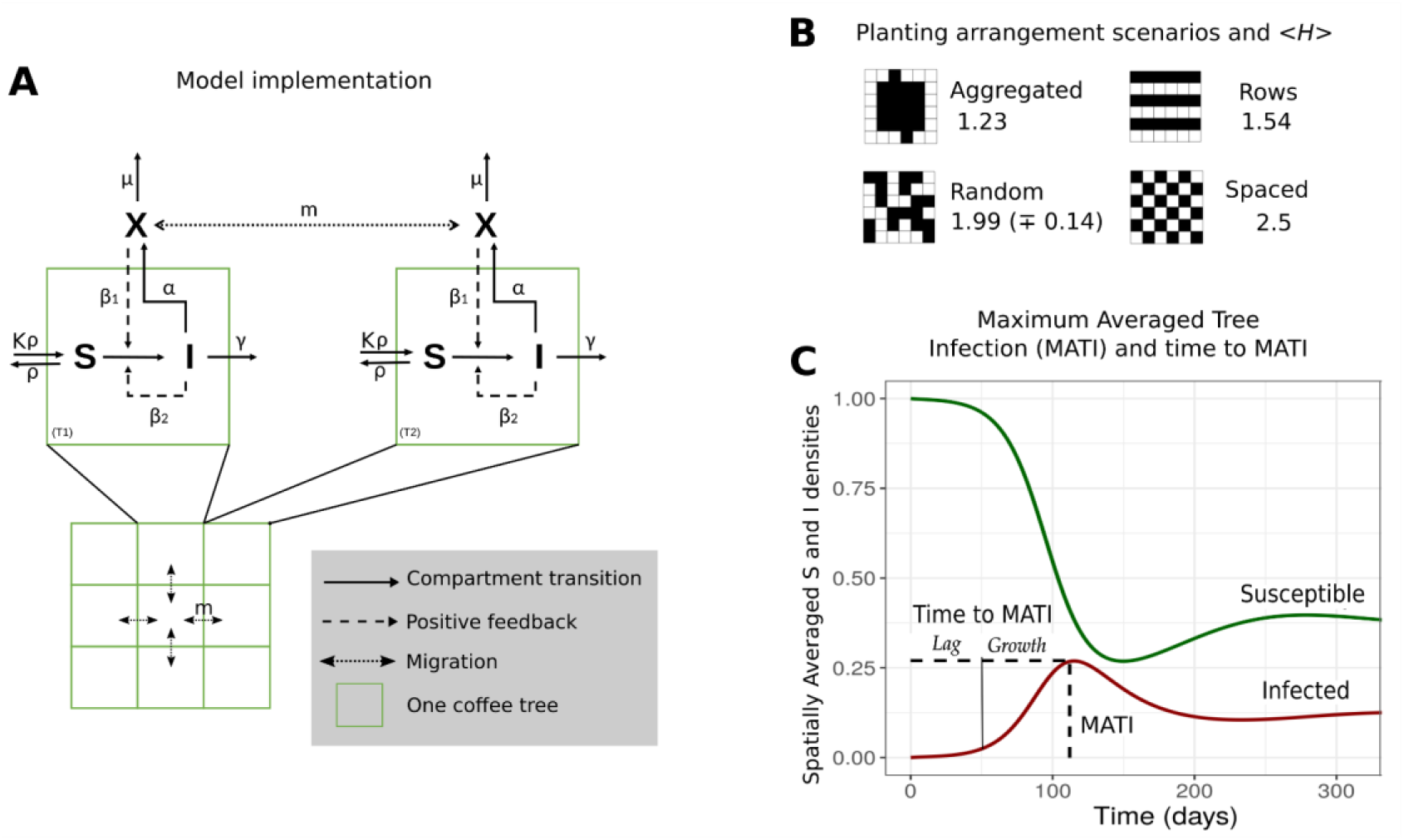
Model Diagram and the different planting patterns considered. A. Model scheme; ***S***: Susceptible leaves, ***I***: Infected Leaves, ***X***: External spores. B. Different planting arrangements and their distance-between-plants index *<H>*. The black squares indicate the presence of a tree and the number is the *<H>* value for that pattern. All the arrangements contain 50 trees in a 10 × 10 lattice (here we depict a 6×6 lattice with the same planting density only for visualisation purposes). C. Basic dynamics for ***S*** and ***I***, Maximum Average Tree Infection (MATI) and time to MATI (divided in two parts: the time lag (*Lag*) and the growth *per se* (*Growth*)). ***X*** follows dynamics similar to ***I*** (see Fig S1.1). Here *γ*=0.056, *α*=0.65, *β*_*1*_= *β*_*2*_= 0.035, *ρ*= 0.011 and *μ*= 0.2 (estimated parameters shown in Table 1).

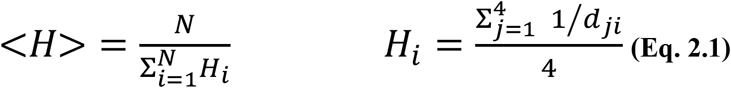

**Table 1.** **Estimated parameter ranges, definition, estimation methods and used values for SIX model simulations**. The letters indicate the direct reference used: a- Rakocevic & Takeshi (2018) b- Mulinge & Griffiths, E., (1974), c- Firman and Wallis (1965), d- Bock (1962), e- Leguizamón-Caycedo et al. (1998), f-Gagliardi et al. (2020), g- Boudrot et al. (2016), h- Rayner (1961), i- Silva-Acuña et al. (1999), j- Nutman, et al. (1963), k- Deepak, K., et al. (2012). GDD: Growing Degree Days. N_sp_: number of infective packages of external spores, N_leaf_: number of susceptible leaves, Ninf: number of infected leaves

| Estimated parameters |  |  |  |  |  |  |  |
| --- | --- | --- | --- | --- | --- | --- | --- |
| Parameter | Definition | Inverse of the time it takes: | Estimation Method | Explanation of the estimation | Units | Range | Used values for simulations |
| $\rho$ | Natural leaf fall rate | One newly mature susceptible leaf (S) to fall | In field measurement (a)<br>Data-Based Estimation (b, c) | Time reported in the field as GDD (a). We also fitted data of leaf development in trees, where coffee rust infection was negligible, to a basic leaf growth equation (b, c). | 1/t | [0.009-0.013] | [0.011] |
| $\beta_1$ or $\beta_2$ | Primary and secondary infection rate | One susceptible leaf (S), in contact with one external package of infective spores (X) or with one infected leaf (I), to become infected | In field measurement (d, e) | Time well documented in several field studies (d, e). | $N_{inf}/(N_{sp} t)$ and 1/t | [0.03-0.04] | [0.035] |
| $\alpha$ | Recruitment rate | One package of spores (X) to leave the plant system from one infected leaf (I) | Data-Based Estimation (f, g, d, h, i) | Estimation based on reports on the number of spores produced per infected leaf and per unit of time, as well as on the relationship between the number of spores in a leaf and the number of spores surrounding the plant system (f, g, d, h, i) | $N_{sp}/(N_{inf} t)$ | [0.1-1.2] | [0.65] |
| $\mu$ | Spore death rate | A new spore to become non-viable. | Data-Based Estimation (j, k) | The spore viability through time has been studied (j, k). We used their data to fit our model. | 1/t | [0.1-0.3] | [0.2] |
| $\gamma$ | Infected leaves fall rate ($\gamma = \rho + \rho_i$) | A newly infected leaf (I) to fall | Data-Based Estimation (c) | Time is shorter than the natural leaf fall time. As far as we know, the former time is not explicitly reported so we used (c) results on susceptible and infected fallen leaves to estimate the leaf fall differences. | 1/t | $\rho + [0.004]$ | [0.015-0.14] |
| Non-estimated parameters |  |  |  |  |  |  |  |
| K | Carrying capacity of susceptible leaves | NA | NA | NA | $N_{leaf}$ | NA | 1 |
| m | Diffusion rate | One package of spores to be fully transported to a 4N- vicinity neighbour square. | NA | NA | 1/t | NA | [0.001 – 0.1] |

where *N* is the number of plants per plot, and *H*_*i*_ is the average of the inverse distance (number of squares) between plant *i* and its four closest neighbours *j* (*d*_*ji*_) along the horizontal and vertical axis. If there are no neighbours along a particular direction, we set *1/d*_*ji*_ = 0. In the case of the random arrangement, we ran 30 configurations and presented the average. Since we chose the same number of trees for all the scenarios (N=50), *<H>* only depends on the spatial arrangement of plants (Fig. 1B).

Each tree follows the two main phases of the coffee rust life cycle: host-pathogen interaction (invasion) and pathogen dispersal, whose dynamics are schematized in Fig1.A and are described by the following system of coupled differential equations (**Eq. 2.2**):

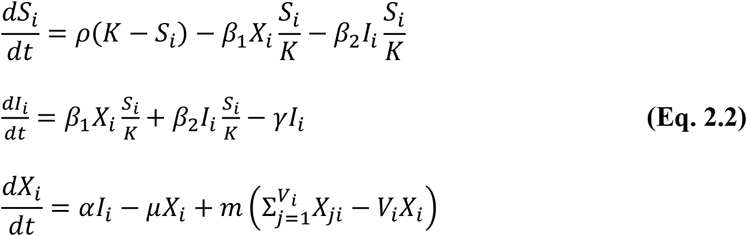

where *Si* and *Ii* are the amounts of susceptible and infected leaves in tree *i. Ii* represents the state where the infected leaf has already produced new infective spores. *Xi* represents the number of infective external uredospores (from now on “external spores”; Supplementary Material 2; Bock, 1962b) in square *i*, with or without a tree. *Xi* does not include spores in the leaves or between them, only the ones that are outside the tree. Host reproduction (*i*.*e*. leaf production and leaf-fall rate of susceptible leaves) is represented by the so-called monomolecular growth (Cunniffe and Gilligan, 2010), where the natural (or non-infected) leaf-fall rate is represented by *ρ* and leaf production rate is equivalent to *Kρ* where *K* is the carrying capacity of susceptible leaves. We took *K* = 1 for simplicity but our results can be rescaled by using reported values for *K* (Burdekin, 1964). This assumes that the new leaf production rate equals the leaf-fall rate of susceptible leaves. The transition from *S* to *I* is subdivided into primary infection arising from the external spores (*X*) and secondary infection occurring by transmission from already infected leaves (*I*). In both cases, the growth in infection is proportional to the fraction of remaining susceptible leaves *S/K*, and to the rates (*β*_*1*_, *β*_*2*_), respectively. Infected leaves can fall and leave the *I* compartment at a rate *ɣ* that must be higher than the non-infected leaf-fall rate (*ρ*). Spores detach from infected leaves and become suspended in the air or fall on the ground, filling the *X* compartment at a rate *α*. External spores die at rate *μ* both in squares with and without trees (black and white cells in Fig. 1B). Finally, *m* is the diffusion rate, which represents the rate at which spores are dispersed to the neighbouring squares. Let us denote as *Vi* the number of immediate neighbours of square *i* (ranging from two to four depending on the location of the square in the lattice), and *Xji* the amount of *X* in the *j*-th neighbour of *i*. Non-directed dispersal is modelled by a diffusion process that takes place from one plant to its four immediate neighbouring squares, mimicking short ranged rust dispersal mediated by splash and plant-to-plant contact in our two-directional planting arrangements. Note that equivalent scenarios could be modelled using an 8 neighbourhood vicinity, but in order to create the spaced arrangement we would have to push trees further away from each other. In empty squares, *S*_*i*_ *= I*_*i*_ *=* 0, so the spores have the following dynamic: 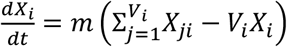. All the parameters are summarised in Table 1.

Our model is a spatially extended case of the SIRX (Susceptible-Infected-Removed-eXternal inoculum) models analysed by Cunniffe and Gilligan (2010) and Gubbins et al. (2000), originally developed for insect-pathogen interactions but widely used for plant-pathogen associations (Swinton and Anderson, 1995). Here we did not include the removed compartment (*R*) since its dynamic is determined by the other compartments and does not impact the whole infection process (this assumption is discussed in the Discussion section). In this sense, our model can be referred to as a SIX model. We assume that infection happens in non-resistant coffee trees with “well-mixed leaves”, when the conditions for the development of coffee rust are optimal (sufficient susceptible leaves and humidity (Nutman et al., 1963)). Factors like plant or rust variability are not explicitly considered, nor the change in abiotic conditions. In this work we do not study the equilibrium points but rather some of the most relevant transient dynamics of coffee rust epidemics like the duration of the growth phase or the maximum infection, as well as the probability of rust invasion in each individual plant (but see (Cunniffe and Gilligan, 2010) for the linear analysis of the model and the supplementary material S1 for the equilibrium points). The invasion criteria is summarised by the parameter *R*_*0*_ which is defined as:

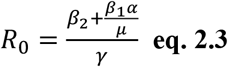

If *R*_*0*_ is less than 1, the single plant system always reaches a-non infective equilibrium. If *R*_*0*_ ≥ 1, *I* and *X* “invade” and S decreases (see Fig. S1.1 and (Cunniffe and Gilligan, 2010) for the main mathematical results on this model).

### 2.3 Plant level parameterization and simulation conditions

We selected the infected leaf-fall rate (*ɣ*) as the sole plant-level variable and fixed the other parameters of the SIX equation (primary and secondary infection, recruitment, leaf-growth, and spore-death rates). We chose the infected leaf-fall rate as it is a variable that can be directly modified by human management (e.g. removing and bagging away the infected leaves). The other parameters such as the infection rates are affected by multiple environmental, genetic and management processes and are more difficult to control. Besides, this decision reduces our computational explorations and simplifies the interpretation of the results. To choose the values of the plant-level parameters correctly and work with meaningful timescales, we estimated the range of each plant or spore parameter from reported data (Table 1) (Bock, 1962a; Boudrot et al., 2016; Deepak et al., 2012; Firman and Wallis, 1965; Gagliardi et al., 2020; Leguizamón-Caycedo et al., 1998; Mulinge and Griffiths, 1974; Nutman et al., 1963; Rakocevic and Takeshi Matsunaga, 2018; Rayner, 1961; Silva-Acuña et al., 1999). We then set each parameter to the mean value for our general simulations, but we included a sensitivity analysis of our results, using the lowest and highest values of the estimated ranges. The infected leaf-fall rate (*ɣ*) was varied above its estimated mean to account for leaf-removal practices that shorten the time for a rusted leaf to fall (*e*.*g*. selective pruning), but we restricted its maximum value to simulate scenarios where rust invades the system (this is, where *R*_*0*_ ≥ 1 considering the other estimated parameters; equation 2.3). The detailed methods for the estimation of each parameter are available in the Supplementary Material S3. To summarise, the model parameters can be estimated from the characteristic time-scale (in days) of a given known process. We first looked for studies that reported those times directly or indirectly. If these data were not available, we fitted specific time-series of leaf infection to linear or exponential models (Table 1).

### 2.4. Simulations and measured variables: MATI and time to MATI

With the chosen parameters, we explored the effect of the leaf-fall rate of infected leaves (*ɣ*), the initial proportion of infected leaves per tree (*I*_*0*_), the planting patterns, and the different plant-to-plant diffusion rates (*m*) on the maximum averaged tree infection (henceforth; MATI) and the time to reach this maximum (in days) (Fig. 1C). The MATI is obtained by averaging tree infection over all trees in the plot for every time step, and then by identifying its overall maximum. This is a common indicator to measure rust infections in the field. The time to MATI was divided into a time lag and a growth period *per se*, following the same criteria used in real time series (see section 2.1 and Fig.1C). The results were then compared with the magnitudes measured in section 2.1 and analysed in each combination of scenarios.

#### 2.4.1. One-plant level simulations

We first ran simulations of the SIX model at the single-tree level (N=1, *m=0* and *i=1*), studying the effect of both the *ɣ* and *I*_*0*_ on the MATI and the time to MATI. In this scenario, the MATI is the maximum local tree infection. We varied the leaf-fall rate of infected leaves (*ɣ*) from 0.015 to 0.14 and the initial proportion of infected leaves per tree (*I*_*0*_) from 0.001 to 0.1 (72 scenarios in total). Each simulation started with *I*_*0*_ infected leaves (*I=I*_*0*_), *1-I*_*0*_ susceptible leaves (*S=1-I*_*0*_), no external uredospores (*X=0*) and ran for 30 000 integration steps using the Euler method (*Δt*=0.01). This method is a standard and common integration procedure for solving discretized partial differential equations in lattices (Koch and Meinhardt, 1994; Elder et al., 1992) and is sufficiently robust and precise for all our modelled scenarios and quantities of interest (with a *Δt* = 0.02, there were no differences in the results within the range of precision used (Fig.S1.6)). This total time represented 300 days, after parameter calibration. The beginning of the simulation corresponds to the start of the optimal conditions for infection (when humidity is sufficient, and leaves are susceptible) and we chose 300 days to be the maximum time for the rust to reach the maximum peak of infection in one year (following the times reported in Fig.2). After this, rust infection is assumed to decrease. This time limit sets a maximum simulation time.

**Fig. 2.**
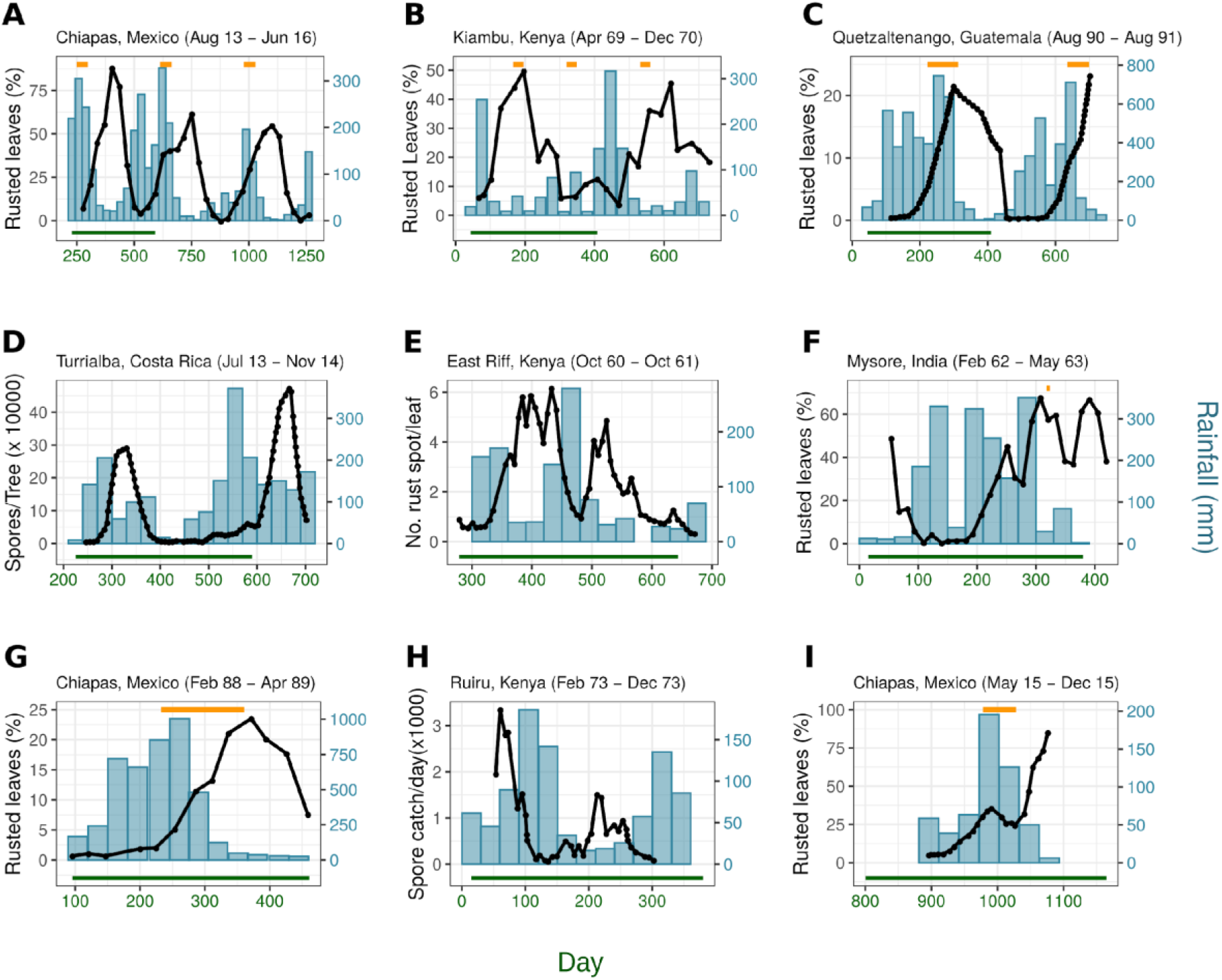
Coffee rust infection dynamics in relation to annual rainfall and harvesting period. The black line represents the amount of rust infection, measured as the average percentage of rusted leaves per tree (A, B, C, F, G, I), or the average number of rust spots per leaf (E), or the number of spores per tree (D), or above the tree (H). The bars represent the monthly rainfall (mm). The orange horizontal segment in figures A, B, C, F, G and I shows the reported harvesting period and the green horizontal line in the bottom of each graph stands for the 365-day period. The plots are ordered from the longest to the shortest period recorded. The references for each plot are: A (Vandermeer et al., 2018), B (Mulinge and Griffiths, 1974), C (Avelino et al., 1993), D (Boudrot et al., 2016), E (Bock, 1962a), F (Ananth, 1969), G (Avelino et al., 1991), H (Becker and Kranz, 1977; Ndolo et al., 2017), I (Vandermeer et al., 2018). For more details, see Supplementary Material 2.

#### 2.4.2 Plot-level simulations

The next step was to run the full model in the 10 × 10 lattice to include the effects of spatial arrangement and diffusion rate on the MATI and time to MATI. We defined scenarios with the four different planting arrangements (aggregated, spaced, random and rows), one initially infected tree in the centre of the lattice ([x,y] =[5,6]), two values of initially infected leaves (*I*_*0*_ =0.001 and *I*_*0*_ = 0.1), and five different levels of diffusion rate (*m*) across three orders of magnitude (ranging from 0.001 to 0.1; or expressed in log for a better visualisation, from −3 to −1). We also chose two of the values of *γ* used in the single plant dynamic (0.015, 0.056) to compare the plant and plot-level results. For the random planting, we took averages over 30 simulations for each combination of the parameters. For each scenario we obtained the MATI and the time to MATI as a function of the diffusion rate, initial infection and *<H>*. We also included the time evolution of the average tree infection with two representative diffusion values (log(*m*)=-3, log(*m*)=-2). As we consider *t* ≤ 300 days, the results of the scenarios can be divided into two cases: **a**. when an infection peak is attained before the 300 days and **b**. when the optimal conditions for rust cease before the peak is reached (creating a maximum at 300 days). Finally, in order to explore the relationship between individual and average dynamics, we registered the values of each tree’s maximum infection and timing and grouped the number of trees that reached a high level of infection (more than 70% of infected leaves) during the same 15-day period. Focusing both on the level of infection of individual trees and their degree of temporal overlapping can shed light on the determinants of the average infection dynamics.

#### 2.4.3. Computational implementation

The model and simulations were implemented in the *Python* 3.7.3 programming language, using the modules *NumPy, SciPy, Pandas, Seaborn* and ran on the LANCIS facilities, at the Ecology Institute of UNAM. The data analyses and figures were done in *Rstudio* 1.2.1335 using *plyr, dplyr, tidyverse, ggplot2* and *patchwork* libraries and *Inkscape 1.0*. All code and data to reproduce results in the work can be accessed at https://github.com/tenayuco/dispersion_plant_arrangement_coffeerust_infection

## 3. Results

### 3.1. Trends in qualitative dynamics coffee rust infection: seasonality and variability in time lags, growth phase and maximum infection

Fig. 2 depicts different coffee rust infection dynamics observed in nine sites, the corresponding rainfall, and, when reported, the harvesting period. Each infection is measured either as the average percentage of rusted leaves per tree or as the average number of spores, during at least one year (green bar at the bottom of each plot). Plotting the time series together enables us to visualise similarities and differences. Firstly, rust infection follows a basic epidemiological cycle consisting of a time lag in relation to the beginning of the rainy season, followed by a growth and a decline phase.

Coffee rust infection also seems to have a rain-forced periodicity: the growth phase always starts after the onset of the rain season (this is clearer for Fig. 2A, B and C where more than one year is reported). This forced periodicity is related to the first phases of the coffee rust infection cycle, where rain is necessary for spores liberation and invasion. The infection reaches a maximum value and declines when the rain season is ongoing (D, H) or has ended (A, C, F, G). In Kenya, where there are two rainy seasons per year, we observe two peaks of infection (Fig. 2B, E and H). The second peaks are substantially lower than the first peaks in East Riff and Ruiru Sites but they are qualitatively relevant to the general dynamics (B, H), as they show that a new infection process began.

The time lag is 82 days on average (ranging from 24.5 (B) to 146 days (C); see Table S1.1). In sites with two rainy seasons, the lag shortens (Fig. 2 B, E, and H). In some cases, coffee rust infection is negligible during the lag (Fig. 2C, D, F and G). The growth phase takes 119 days on average, ranging from 68 (I) to 181 days (C) (Table S1.1). It is worth noting that the coffee rust infection cycle takes about 30 days, therefore rust probably undergoes multiple infection cycles during one epidemiological phase, as discussed in Kushalappa and Eskes (1989). The “time to maximum infection” comprises the time lag and the growth phase and ranges from 95 (I) to 325 days (C). The maximum average tree infection ranges in turn from 20 to 80% of infected leaves per tree. Therefore, the timing and intensity of an infection at a given site are largely variable. Interestingly, the harvesting period correlates with the build-up (growth phase) (Fig. 2A, C, G and I) (see the discussion section). All values for the MATI and time to MATI are summarised in Table S1.1.

### 3.2. The SIX parametrised model reaches a rust-infected equilibrium

We estimated the natural (non-infected) leaf-fall time, primary and secondary infection, and spore death rates directly from reported data (S3; Table 1). The leaf-fall time of susceptible leaves (*1/ρ*) ranges between 74 and 108 days. *β*_*1*_ and *β*_*2*_ can both be interpreted as the inverse of the time taken for one susceptible leaf to become infected, either for being in contact with external spores or with an infected leaf. If we assume that one package of external spores infects one leaf at a time (this is N_sp_/N_inf_ =1) both times are equal and range from 25 to 30 days (*1/β*_*1*_ and *1/β*_*2*_). Finally, the uredospore viability goes from 3 to 10 days (*1/μ*) (Table 1). The time taken for a newly infected leaf to fall (***1/****ɣ*) was estimated indirectly as how much shorter this time was in comparison to *1/ρ*. Infected leaves take on average from 35 to 53 days less than susceptible leaves to fall.

We express *ɣ* as *ρ + ρ*_*I*_, where *ρ*_*I*_ is the increment in leaf-fall rate due to coffee rust infection. The recruitment rate (*α*) range is broad [0.1-1.2] since it was estimated indirectly from several studies (Table 1). We use *ρ* = 0.011, hence the mean value of *ɣ* is equal to 0.015 (Table 1). Given the values chosen for the other parameters here, the invasion criterion can be expressed as *R*_*0*_ = 0.15/*ɣ* ≥ 1 (see equation and Fig.S1.1). In this sense, to simulate rust infected scenarios and to account for leaf-removal practices (see section 2.3), *ɣ* was varied between 0.015 to 0.14 (*R*_*0*_ ∈[1, 10]).

### 3.3 In one-plant simulations, ɣ affects both the MATI and time to MATI, and I_0_ affects the time to MATI through the time lag

Figs. 3A and B display the maximum averaged tree infection (MATI) and the days to MATI, for one-tree simulations (N=1), varying the leaf-fall rate (*ɣ*) and initial proportion of infected leaves (*I*_*0*_). MATI decreases with the infected leaf-fall rate (Fig. 3A and C). When *ɣ* is equal 0.015, the maximum infection is around 75%. When *ɣ* is higher than 0.1, MATI drops to zero. This effect is independent of the number of infected leaves in the systems’ equilibria predicted by *R*_*0*_ (section 2.3; Fig. S1.3; see Fig. S1.1 for the influence of *ɣ* on equilibria and stability). *I*_*0*_ does not affect the MATI whatsoever (Fig. 3D). Time to MATI is affected by both *ɣ* (Fig. 3B and C) and *I*_*0*_ (Fig. 3B and D). In general, when *ɣ* increases, the time to reach MATI increases because the curve flattens (Fig. 3C), affecting the growth phase duration but not necessarily the time lag (see Fig. 2). On the other hand, when *I*_*0*_ is low, the time to MATI increases without changing the MATI (Fig. 3B and D). In other terms, *I*_*0*_ affects the time lag but not the growth phase *per se* (Fig. 3D). When MATI drops to 0 (with high values of *ɣ)* the time to MATI is not relevant since the maximum infection can be reached at the beginning of the simulation.

**Fig. 3.**
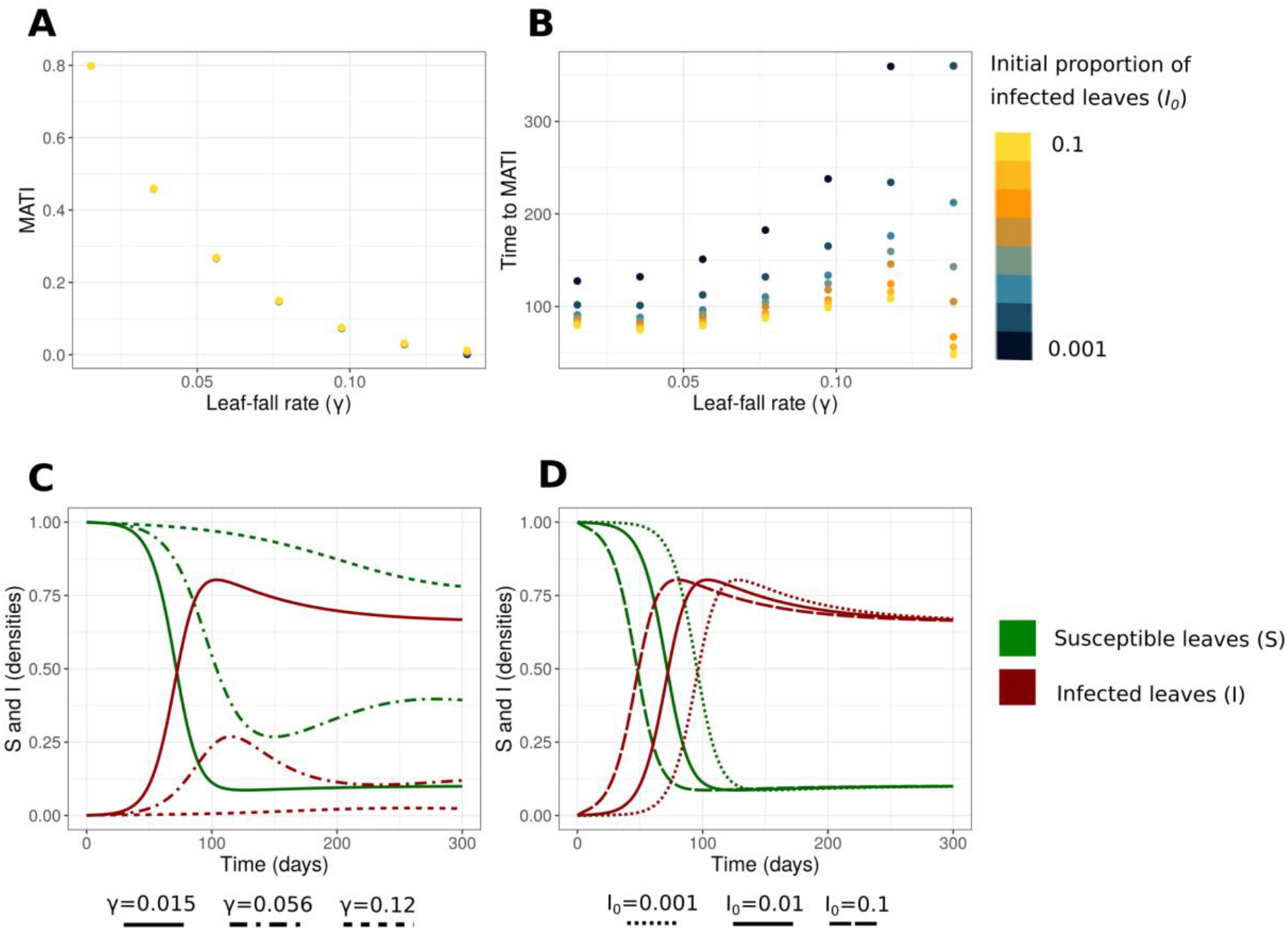
MATI and time to MATI in isolated coffee plants (N=1). A, B: MATI and days to MATI in function of the leaf-fall rate of infected leaves (*γ*) and initial proportion of infected Leaves (*I*_*0*_). C: Example of infection dynamics for susceptible and infected leaves (see Fig. S1.2 for the evolution of the spores ***X***) with different values of *γ* and a fixed *I*_*0*_ = 0.01. D: Example of infection dynamics with different values of *I*_*0*_ and a fixed *γ*=0.015. The values of *α, β*_*1*,_ *β*_*2*_, *ρ* and *μ* are shown in Table 1.

### 3.4. In the spatially explicit model, diffusion rate and the planting arrangement jointly modify MATI and days to MATI in a threshold-dependent manner and modify the curves of infection, while I_0_ modulates the time lag and γ decreases the MATI in all scenarios

We plotted in Fig.4 the MATI and time to MATI for each combination of planting arrangement, diffusion, and initial conditions (*I*_*0*_), and in Fig. 5 the time evolution of the average tree infection with two representative diffusion values (log(*m*)=-3, log(*m*)=-2). We used two values of *γ* (0.015, 0.056) but as the results were qualitatively similar, we only present the plots here for *γ*=0.015 (but see Fig.S1.7 for the case of *γ*=0.056). As we explained in the methods sections, results are divided into two cases: **a**. when an infection peak is attained before the 300 days and **b**. when the optimal conditions for rust cease before the peak is reached (creating a maximum at 300 days).

The variations of the time to MATI and MATI for all the combinations of scenarios fall within the range reviewed in section 3.1 (see Fig.2 and Tables S1.1 and S1.2). In all the spatial arrangements, when diffusion rate increases, MATI increases, and days to MATI decrease (Fig. 4).

**Fig. 4.**
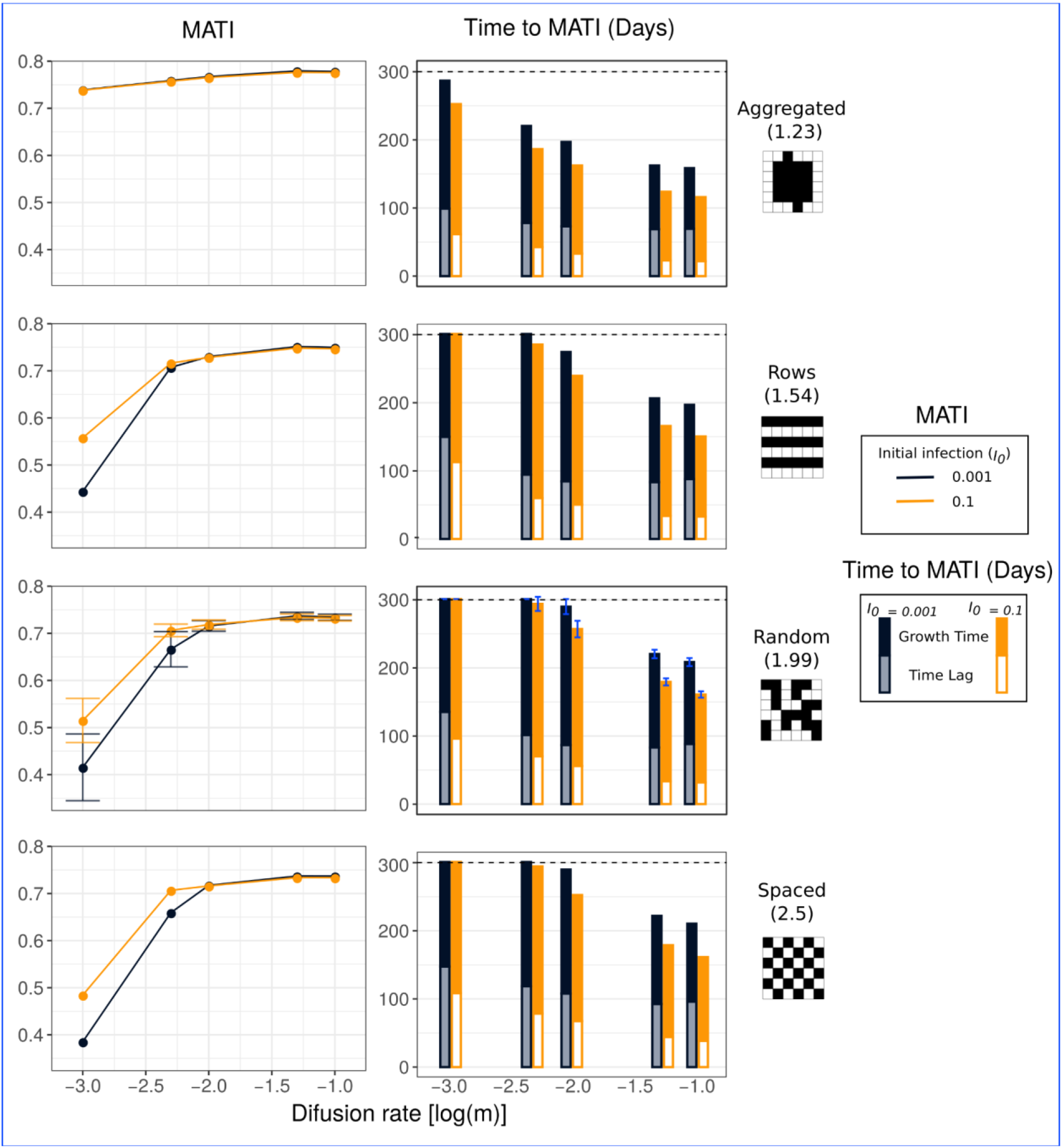
MATI and time to MATI for all simulations. Simulations of four different planting arrangements (aggregated, random, rows and spaced, with their respective distance-between-plants index *<H>*) with five levels of diffusion rate (log(*m*) between −3 and −1) and two levels of *I*_*0*_ (0.001 and 0.1, dark and orange lines respectively). The bars representing the time to MATI are divided into the time lag and the growth period *per se* (light blue vs black and light orange vs dark orange). The dotted line represents the 300 days limit of the simulations. For the random arrangement, each point/bar represents a 30-simulation average, and the error bars the standard deviation. The values of *α, β*_*1*,_ *β*_*2*_, *ρ* and *μ* are shown in Table 1. We used *γ*=0.015 (see Fig.S1.7 for *γ*= 0.056).

In the aggregated planting arrangement MATI is always reached before 300 days (ranging from 116 to 287 days) and has values above 70% of infected leaves, only slightly modified by the diffusion rate. In the rows, random and spaced arrangement the change of diffusion rate has a more pronounced effect on the MATI. When the diffusion rate is minimal (log(*m*) = −3) the time to MATI is 300 days (case **b; Fig. 4 and Fig. 5**). In those scenarios, MATI reaches values that range between 35% and 55% of infected leaves, depending on how advanced the infection was at 300 days. If the diffusion rate is larger (more than −2.5 or −2 depending on the arrangement and the initial infection), the time to MATI becomes less than 300 days for these three arrangements (varying between 151 and 294 days). In these scenarios, an infection peak is reached before the end of optimal conditions (*i.e* before the end of the simulation; case **a**).

**Fig. 5.**
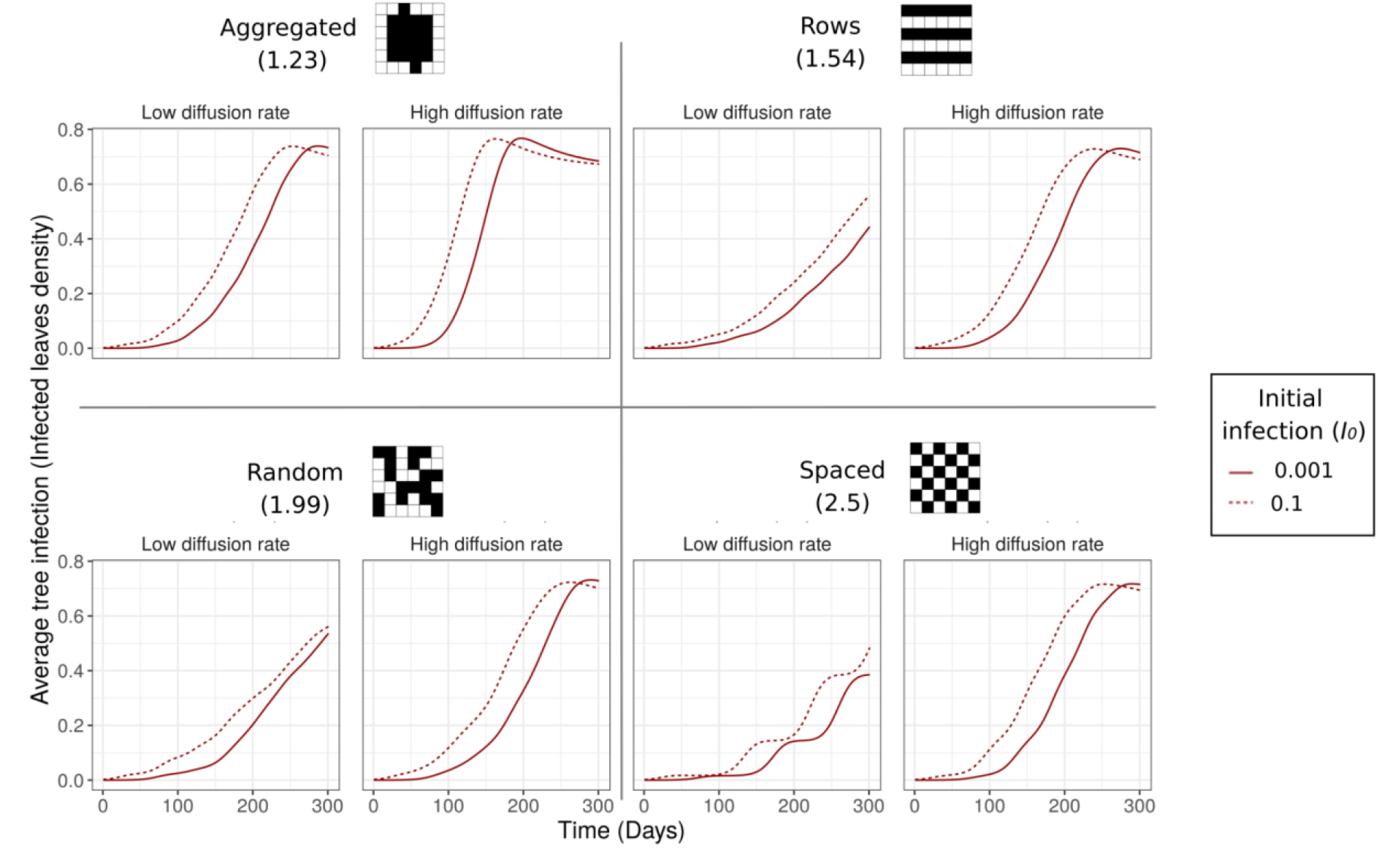
Examples of average tree infection dynamics for plants in a plot. Simulations of four different planting arrangements (aggregated, random, rows and spaced) with two levels of diffusion (low: log(*m*)=-3, high: log(*m*) = −2) and two levels of *I*_*0*_ (0.001 and 0.1). The lines represent averages over all the trees of the plot at a given time. The values of *α, β*_*1*,_ *β*_*2*_, *ρ* and *μ* are shown in Table 1. We used *γ*=0.015.

In other terms, for lower values of diffusion rate (log(*m*) = −3) the change of the diffusion rate affects the MATI according to a critical distance between plants (*<H>*): when *<H>* is higher than 1.5, this change greatly modifies the MATI (Fig.4 and Fig. S1.4), when *<H>* is lower than this threshold, the MATI is slightly affected. For higher values of the diffusion rate (log(*m*) > −3), the differences in aggregation become less relevant. Interestingly, in all scenarios, the diffusion rate modifies the growth time *per se* (that varies between 151 and 205 for log(*m*) = −3, and 89 and 129 for log(*m*) = −1) (see Fig. 4 and Table S1.2). For low diffusion rates, this results in flatter curves leading to lower MATI (Fig. 5). Moreover, differences in planting arrangements do modify the qualitative average tree infection dynamics, creating smoother or rougher curves (Fig. 5). In particular, the spaced arrangement combined with a low diffusion rate, generates multiple steps in the epidemiological dynamics (Fig. 5). This is similar to what we observe in Fig.2 B or H. Random and row planting arrangements did not present any relevant differences between them.

*I*_*0*_ mainly affects the time lag (that varies between 70 and 150 days when *I*_*0*_=0.001 and between 22 and 113 days when *I*_*0*_=0.1) (see Fig. 4 and Table S1.2). The curves of Fig. 5 are thus shifted along the time axis as *I*_*0*_ varies. It is noteworthy that the time lag can be higher than in the one-plant simulation scenarios (that go from 10 to 50 days as *I*_*0*_ is varied). This increase in the time lag can lead to a lower MATI if the curve of infection does not reach a maximum peak and is still increasing at the end of the simulation (Fig.4 and Fig.5).

All of our results were qualitatively robust to the variation of the other plant-level parameters within their estimated range, except for the lowest value of α (α=0.1) that significantly decreases the MATI in all scenarios (Fig.S1.7). Finally, as in the plant-level simulation, the MATI decreases and the time to MATI increases when the infected leaf-fall rate (γ) increases (Fig.S1.7.B), especially for values greater than the estimated range (γ = 0.056).

### 3.5. The MATI and time to MATI are determined by the number of individual plants that reach a high level of infection before the end of the simulation, and their degree of temporal overlapping

The level of infection of individual trees and their degree of temporal overlapping determine the average infection dynamics. Both variables are affected by the diffusion rate and the planting arrangement. We represent in Fig. 6 the proportion of trees that reached a high level of infection (>70% of infected leaves) during the same 15-day period, and the accumulated proportion of corresponding highly infected trees (dotted line), which indicates how much the epidemic has expanded in space up to a given time (see S4 for a full plot visualisation and refer to Fig. S1.5 for the values of each tree’s maximum infection and timing).

**Fig. 6.**
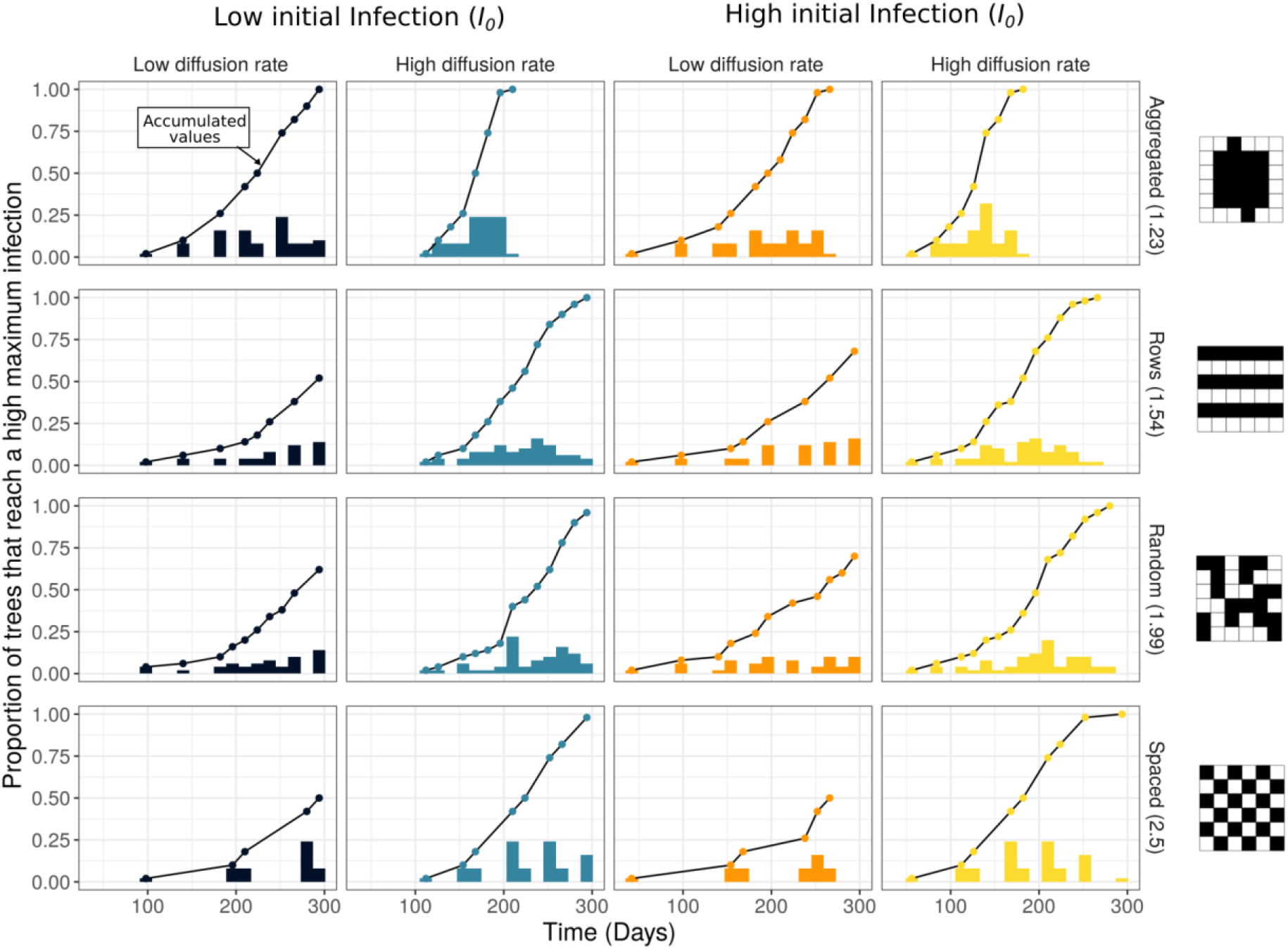
Proportion of trees that reach a high maximum infection (>0.7 of infected leaves) during the same 15-day period. Histograms for the different planting arrangements (aggregated, random, rows and spaced) with two levels of diffusion (low: log(*m*)=-3, high: log(*m*) = −2) and two levels of *I*_*0*_ (0.001, 0.1; dark/blue and orange/yellow colours respectively). The black dotted line represents the corresponding accumulated proportion of trees in the plot that reach a high maximum infection. The values of *α, β*_*1*,_ *β*_*2*_, *ρ* and *μ* are shown in Table 1. We used *γ*=0.015.

Planting arrangement substantially influences the overlap between individual tree dynamics. This explains the different shapes of the average infection curves (Fig.5) as well as the changes in MATI and time to MATI. When plants are spaced, in rows, or randomly distributed, the distribution of the number of highly infected trees over time is broader than in the aggregated pattern, especially when diffusion is high (Fig. 6; yellow and light blue bars). This means that in the first three scenarios the temporal overlap of highly infected trees decreases, which causes a lower MATI compared to the aggregated pattern (Fig. 4 and Fig. 6).

The diffusion rate, combined with the planting pattern, affects both the overlap and the individual times at which the maximum individual infection is reached. When diffusion is low (dark blue vs light blue, and orange vs yellow bars in Fig. 6) the distribution is spread out and slightly shifted to the right (see also the accumulated curve). At low diffusion, for the spaced, rows and random patterns, only 50 to 75% of the trees reach the 70% of infected leaves threshold before the end of the simulation. The other trees with low levels of infection will contribute little to the average tree infection value, lowering the MATI (Fig.4 and 5 and Fig.S1.5). In the aggregated pattern or when diffusion is high, all the trees reach 70% of infected leaves (the accumulated curves sum up to 1). Interestingly, high values of *<H>* (which characterises rows, random and spaced arrangements) combined with a low diffusion rate (dark and orange bars) generate isolated outbreaks of highly infected trees (Fig.6). In particular, in the spaced pattern, individual trees reach high infections at well separated times, explaining the multi-stepped dynamics of the spatially averaged prevalence in Fig. 5. Fig. 6 also shows that a lower amount of initially infected leaves (*I*_*0*_) delays the initial infection without significantly changing the dynamics.

## 4. Discussion

One of the central issues in coffee rust epidemiology is to understand the factors that explain the great variability in the magnitude of the infection peaks and their timing, both at the plant and plot levels (Avelino et al., 1991; Li et al., 2022; Rosas et al., 2021). To address this issue, many studies have focused on the microclimate (Avelino et al., 2006), plant resistance (Talhinhas et al., 2017), or the susceptibility of leaves during specific stages of the coffee crop cycle (Salgado et al., 2008). Nevertheless, none of these approaches has succeeded to fully explain the differences in timing and intensity of the epidemics occurring in plots from the same site with equivalent meteorological and plant genetic conditions. Here we hypothesised that spatial patterns created by rust dispersal in different planting arrangements might be affecting both variables.

We first documented known trends for the timing and magnitude of the epidemic peak, as well as their variability, by comparing epidemiological curves from different sites (Fig. 2). From this analysis we distinguish two phases in the epidemic timeline: a time lag and a growth phase *per se* (Fig. 2 and Table S1.1). We also noted that the harvesting period correlated with the build-up or growth phase (Fig. 2A, C, G and I). This is consistent with harvesting and rust infection being related to fruit development and fruit load (Avelino et al., 1993; Motisi et al., 2022; Salgado et al., 2008). Nevertheless, the movements of workers during harvesting might also promote rust dispersal and reinforce the infection (Motisi et al., 2022). Our parameterised SIX model recreates the coffee rust epidemic and helps us to explore some of the mechanisms that may determine properties of the maximum peak and its timing (Fig.3-6). The values of the MATI and time to MATI were within the range reported in field studies. This qualitatively validates the estimates of the model parameters, and supports that different planting arrangements and different diffusion rates might provide possible explanations to the variation observed in real dynamics.

In particular, we showed that both the planting arrangement and the diffusion rate jointly modify the MATI and time to MATI by preventing some plants from reaching their maximum peak (Fig.4, Fig. 5, Fig. 6), thus explaining part of the variability observed in plots with otherwise similar conditions. In particular the aggregated spatial arrangement highly increases the MATI compared to the other planting arrangements that did not present quantitative differences between them (see videos in S4). The effects of the planting arrangement are noticeable in the different curves of Fig. 5, which can be multi-stepped, or smooth (like some of the curves from real data presented in Fig.2). The multi-stepped pattern is associated with a higher distance between plants, in particular in the spaced pattern, where plants have no direct neighbours. In this sense, the differences of the effect of planting patterns on the MATI and curves of infection were more related with a planting aggregation threshold (or a minimal distance between plants) than with the spatial arrangements *per se*. Moreover, a low *I*_*0*_ increases the time it takes to reach the maximum peak of infection at the plant level and plot level, due to delays in the start of the growth phase (Fig. 4, 5). The time lag differences reported in sites with similar phenological timing (Li et al., 2022) can thus be explained by differences in the initial infection and planting arrangements, and not only by the positive correlation between fruit charge and plant susceptibility to infection (Avelino et al., 1993; Salgado et al., 2008). Finally, we found that high infected leaf-fall rate (*ɣ*) strongly affects the whole-plot infection dynamics and we hypothesise that this quantity is a key aspect for coffee-rust monitoring or intervention at the plot scale (Park et al., 2001).

These results have a practical significance regarding coffee rust management. They highlight the importance of reducing rust dispersal rates, either by having other trees in the plot or by planting coffee plants at larger distances (independently of the spatial arrangement *per se*) in order to uncouple individual dynamics and reduce plot infection. In particular, one should avoid the aggregated arrangement, this is, *<H>* values larger than 1.5 (Fig. S1.4; videos S4). This result complements those of Hajian-Forooshani and Vandermeer (2021), who tested different critical dispersal distances between coffee plants and reported that the regularity of the planting arrangement pattern modulates the time to reach fully rust-infected coffee plots. This is relevant for farmers, who balance their planting density according to their production necessities and the risk of plot epidemic (Ehrenbergerová et al., 2018). In general, conventional coffee management produces very densely aggregated planting patterns that might be prone to coffee-leaf rust invasion, contrarily to more rustic or ecological coffee plantations that intercalate different kinds of trees between the coffee plants (Moguel and Toledo, 1999). Besides, Boudrot et al. (2016) and Gagliardi et al., (2020) reported that the functional traits of plants related to shade (foliage density, shade percentage) modify the relative importance of wind and rainfall in uredospore dispersal across the plot. Understanding the role of the initial infection and dispersal rate on the time lag is also relevant for rust-control procedures that rely heavily on the timing of the epidemic, such as pruning, management of shade (Boudrot et al., 2016; Liebig et al., 2019; Soto-Pinto et al., 2000), or fungicide application (Burdekin, 1964; Mulinge and Griffiths, 1974). Additionally, the leaf-fall rate of infected leaves can also be modified with management practices but may have an ambiguous effect on the maximum infection: on the one hand, farmers could decrease the maximum infection by removing and bagging away the infected leaves continuously (thus decreasing the source of new reinfections from infected leaves either on the ground or in the plant system); but on the other hand, due to the farmers’ movement between the trees, removing the leaves could trigger the dispersal of uredospores by contact and further propagate the epidemic.

The present model also allows us to make recommendations on the ways coffee rust prevalence can be measured and monitored. Here we worked with the average tree infection because it is one of the more common ways of measuring coffee rust prevalence in coffee plots (Fig. 2 and S2). Nevertheless, this variable is highly dependent on the progression of the infection in the chosen plants and their degree of temporal overlap due to spatial factors (Fig. 6). Additionally, the average over trees can be an inappropriate measure for the plot infection when individual infections follow a non-normal distribution that changes during the epidemic. Even if we supposed the infection to follow a normal distribution, the mean without the variance would not be sufficient. Other ways of measuring plot infection in the field could consist in drawing spatially explicit plot infection maps (Li et al., 2022; Rosas et al., 2021; Vandermeer et al., 2018).

In our model, we considered rust dispersal from one plant to its four closest neighbours through splash and leaf-to-leaf contact. This assumption was linked to the spatial arrangements that were only significantly different in the 4-neighbour vicinity. However, in future work it will be important to consider other mechanisms and models for spore dispersal that consider an 8-neighbour range or long-range dispersal. On the one hand, it is known that changing the neighbourhood of interaction can change the probability of plot invasion (Park et al., 2001). Also, having a different dispersal rate depending on whether the plants are in contact or not, can lead to critical transitions in infection dynamics (Hajian-Forooshani and Vandermeer, 2021; Vandermeer et al., 2018). On the other hand, if we were to consider other dispersal mechanisms such as wind gusts or human action during harvesting (Becker and Kranz, 1977), the planting arrangements might play a less important role in plot infection. This is because spores would reach farther trees and attenuate the relevance of the plot geometry. In those scenarios, the amount of shade-trees that can act as wind barriers would be crucial to reduce dispersal (Gagliardi et al., 2020) and uncouple the individual tree infection dynamics. Moreover, the model considers only one initially infected tree located in the centre of the plot. Other choices should not qualitatively change our results, but might have an effect on the general times of infection. The effect of multiple sources of infection should also be studied in future works. It is also important to keep in mind that our model assumes that infected fallen leaves are removed from the system (bagged away, for example) and cannot spread infection to neighbouring plants or reinfect. In many sites, these fallen leaves with spores likely contribute in a meaningful way to infecting plants (Zachary Hajian-Forooshani, personal obs). This could be added in the future by defining an additional R compartment in the SIX equation. Nevertheless, as coffee leaf rust is a biotroph, the survival of spores in the fallen leaves is expected to be limited.

Weather covariates like high rainfalls might also influence plant-level parameters such as the recruitment of spores from infected leaves (*α*) by washing off spores from the leaves. This is likely to strongly affect the general dynamics of infection (*e.g*. reducing the maximum infection; Fig.2 and Fig. S1.7; Motisi et al., 2022) and should be considered in future works.

Finally, our model represents coffee rust infection but does not assess coffee production reduction. This relationship is not necessarily linear and must be considered for any practical recommendations (Cerda et al., 2017).

Control and management practices of coffee rust disease have commonly underestimated the role of dispersal and the relationship between plant and plot dynamics. Our work contributes to a better understanding of this relationship and can be used as a null-model to study the role of specific dispersal mechanisms, such as wind, splash or human mobility (see for example, Avelino et al., 2012; Becker and Kranz, 1977; Gagliardi et al., 2020; Hajian-Forooshani and Vandermeer, 2021). It mostly aimed at exploring the general and qualitative mechanisms behind the patterns observed in real systems (Fig.2). Adding context-specific conditions would enable us to do some data-fitting and quantitative comparisons in a more detailed way than the general times of MATI and time to MATI presented here. Recreating and understanding the main mechanisms of coffee rust dispersal is crucial to the development of a preventive approach to epidemics. It informs the management processes that seek to balance the different effects that some practices have on the infection and dispersal stages, in order to design resilient agroecological coffee systems to coffee rust or other epidemics.

## Supporting information

S2_complementaryDataFig2

S1_additionalFiguresTables

S3_parameterEstimation

S4_videoPlotInfection

## 5. Acknowledgements

EMVC is a doctoral student from the Programa de Doctorado en Ciencias Biomédicas, Universidad Nacional Autónoma de México and has received CONACyT scholarship 686776. MB acknowledges financial support from UNAM-DGAPA-PAPIIT (IN207819). CG acknowledged the graduate program Posgrado en Ciencias Biológicas, Universidad Nacional Autónoma de México and CONACyT scholarship 743257. EM thanks Fabian Aguirre for all the discussions and guidance in mathematical analysis and Elisa Lotero for her help and support. The authors thank Rodrigo García Herrera for technical support and two anonymous reviewers for very detailed comments that greatly improved the paper. We also thank members of *La Parcela* Laboratory for their valuable comments and suggestions. This article covers part of the requirements for EM to obtain his PhD Degree in the doctoral program.

