## Supplementary material for "Dispersal and Plant Arrangement Condition the Timing and Magnitude of Coffee Rust Infection": S2_complementaryDataFig2

| Figure | Rust Measurement | Time Range | Rust data and sampling Measurement method |
|---|---|---|---|
| A | Percentage of rusted leaves | AUG 2013- JUN 2016 | All leaves, 744 plants, 6 coffee plants in the center of each of 124 (50 m × 50 m) (1/4 ha) sampling quadrats regular grid within 45 ha plot |
| B | Percentage of rusted leaves | APR 1969- DEC 1970 | Random samples of seventy leaves were collected from each of the nine central trees in each sub-plot |
| C | Percentage of rusted leaves | AUG 1990- AUG 1991 | 6 branches (2 low, 2 middle, 2 top) in 50 trees in 2 plots (25 per plot) of 0.7 ha each |
| D | Total number of spores | JUL 2013- NOV 2014 | Inoculum quantity was assessed every 21 days in (6 branches, at each level of the plant, of 6 different plants in a 775m2 plot) |
| E | No. rust spot/leaf | OCT 1960- OCT 1961 | 100 leaves were collected at random from each plot, an average of five from each tree, and the numbers of individual rust spots counted on each leaf. These were taken from the lower half of the tree, where maximum infection usually occurs |
| F | Average number of lesions per leaf | FEB 1962- MAY 1963. | Ten leaves were collected from the first tree in each plot and six from each of the other trees, to give a total of 100 leaves from each plot. Young leaves were avoided and leaves were collected from all parts of the bush to get representative samples. |
| G | Percentage of rusted leaves | FEB 1988- APR 1989 | ND |
| H | Average weekly spore catch/day | FEB 1973- DEC 1973 | Inoculum was assesed with an air Burkard Trap with a suction capacity of 600 l air / h, the spore capture values represent a sample of 0.6 * 24 = 14.4 m3 air. A 1.25 m |
| I | Percentage of rusted leaves | MAY 2015- DEC 2015 | Three 10 × 10 m plots were set up in three distinct areas. All leaves, all susceptible coffee plants. There are also non susceptible plants that were ignored. |

| Sample frecuency (mean) | Sample size | Coffee variety |
|---|---|---|
| Monthly | Uniform sample (n=744, 6 plants in 124 sampling quadrats of 50*50 m) among approximatively 51 150 plants (31 ha) | Coffea arabica -rust susceptible |
| Monthly | Uniform Sample (n=9) among approximatively 50 trees | Coffea arabica |
| Monthly | Random sample (n=25) among approximatively 3500 plants | Dwarf varieties- Red Catuai |
| Every 21 days | Random sample (n=6) among approximatively 400 plants | Dwarf cultivar Caturra - Coffea arabica |
| Weekly | All trees were sampled on a 20 tree plot (well mixed) | ND |
| Every 2 weeks | All trees were sampled on a 16 tree plot (well mixed) | Coffea arabica (Kents) |
| Monthly | ND | Coffea arabica - Dwarf varieties |
| Every day | 1 sample in an experimental plot | Coffea arabica - Frenc Mission |
| Every 8 days | All trees were sampled on a 165 tree plot (well mixed) | Coffea arabica -rust susceptible |

| Managment characteristics | | |
|---|---|---|
| Plot Density | Shade | Fungicides |
| 0.9 m between individuals within a row and 2 m between rows.Assuming 1m2 space ocuppied by plant, we get 33*50 = 1650 trees/ha | Shade uniformy distributed (Vandermeer et al. 2008). The dominant shade trees are comprised of several Inga species, Alchornea latifolia (Swartz) and Trema micrantha (Blume) | ND |
| Experimental plot: 50-tree. Density non specified | ND | Unsprayed control |
| High density plot (5000 plants/ha). | Inga spp. | Yes. In only 1 plot: 4 to 5 application cyproconazole (10%) at 0.06 % |
| High density plot (5000 plants/ha, 2*1 m) | Average shade cover was 66 and 64% (Lemon, 1967). Trees: Erythrina poeppigiana, Chloroleucon eurycyclum | Copper-based fungicide and cyproconazole. 2 application per year |
| Experimental plot: 20-tree. Density non specified | Guard trees surrounding block | Unsprayed control |
| Experimental plot: 16 trees. Density non specified | ND | Unsprayed control |
| High density plots (density non specified) | ND | ND |
| Dense shade, so probably low density | Extensive managment | ND |
| 0.9 m between individuals within a row and 2 m between rows.Assuming 1m2 space ocuppied by plant, we get 33*50 = 1650 trees/ha | Shade uniformy distributed (Vandermeer et al. 2008). The dominant shade trees are comprised of several Inga species, Alchornea latifolia (Swartz) and Trema micrantha (Blume) | ND |

| Climatic conditions | | | |
|---|---|---|---|
| Site | Annual rainfall and other characteristics | Mean altitude (m a.s.l.) | Mean Temperature |
| Soconusco Region of southwestern Chiapas (15° 10′ N, 92° 20′ W) | 4500 mm | 1090 | 31 ° C |
| Deepdene (Coffee State), Kiambu, Kenya | 1140 mm | 1700 | 18.8 °C |
| Libertad Plantion, Sierra Madre, Colomba, Quetzaltenango, South-East Guatemala | 3000 mm | 720 | ND |
| Experimental station (CATIE), located in Turrialba, Costa Rica (9°539N, 83°389W). | 2,996mm. No marked dry season. However, rainfall is less abundant between February and April | 600 | 22°C |
| Four East Rift-Kenya sites | 797 mm. Rainfall is biannual and ummer have more rainfall | 1738 | ND |
| Coorg District Mysore State, India | 2783 mm | 900 | 21°C |
| Tapachula, Chiapas | 2600 mm | 730 | 23.5 °c |
| Kenya Coffee Research Station in Ruiru (36 ° 54 'East Longitude and 1 ° 5' South Latitude) | 797 mm. Rainfall is biannual and ummer have more rainfall | 1585 | 19.5 °C |
| Soconusco Region of southwestern Chiapas (15° 10′ N, 92° 20′ W) | 4500 mm | 1090 | 31 ° C |

| Rain time series | | Harvesting | |
|---|---|---|---|
| **Source** | **Measurement** | **Reported** | **Range** |
| World weather online-Soconusco | Monthly | Yes | 258- 288 (15 Sep- 15 Oct) |
| Same site as rust | Monthly | Yes | 166-196 (15 Jun-15 Jul), 319-349 (15 Nov-15 Dec) |
| Same site as rust | Monthly | Yes | 222-312 (10 Aug- 8 Nov) |
| Same site as rust | Daily | Yes. Time range non specified | NA |
| Same site as rust | Daily | ND | ND |
| Same site as rust | Daily | Yes | 321 |
| Same site as rust | Monthly | Yes | 233-361 (1 Sep- 27 Dec) |
| Ruiru Kenya for averages. Then Nairobi for time series | Monthly | ND | ND |
| World weather online-Soconusco | Monthly | Yes | 258- 288 (15 Sep- 15 Oct) |

| Reference (to see full reference, see main text) |
| --- |
| *Vandermeer, J., Hajian-Forooshani, Z., & Perfecto, I. (2018). The dynamics of the coffee rust disease: an epidemiological approach usin* |
| *Mulinge, S. K., & Griffiths, E. (1974). Effects of fungicides on leaf rust, berry disease, foliation and yield of coffee. Transactions of the Br* |
| *Avelino, J., Toledo, J. C., & Medina, B. (1993). Développement de la rouille orangée (Hemileia vastatrix) dans une plantation du sud-ouest du Guatemala et évaluation des dégâts qu'elle provoque. ASIC 15th International Conference on Coffee Science, 293–302.* |
| *Boudrot, A., Pico, J., Merle, I., Granados, E., Vílchez, S., Tixier, P., … Avelino, J. (2016). Shade effects on the dispersal of airborne Hemileia* |
| *Bock, K. R. (1962). Seasonal periodicity of coffee leaf rust and factors affecting the severity of outbreaks in Kenya Colony. Transactions* |
| *Ananth, K. C. (1969). Timing and frequency of spraying for control of coffee leaf rust in southern india. Experimental Agriculture, 5(2)* |
| *Avelino, J., Muller, R. A., Cilas, C., & Velasco, P. H. (1991). Development and behaviour of coffee orange rust (Hemileia vastatrix Berk. and Br.) in plantations undergoing modernization, planted with dwarf varieties in South-East Mexico. Café Cacao Thé.* |
| *Becker, S., & Kranz, J. (1977). Vergleichende Untersuchungen zur Verbreitung von Hemileia vastatrix in Kenia/Comparative studies on* |
| *Vandermeer, J., Hajian-Forooshani, Z., & Perfecto, I. (2018). The dynamics of the coffee rust disease: an epidemiological approach usin* |
