## Supplementary material for "Dispersal and Plant Arrangement Condition the Timing and Magnitude of Coffee Rust Infection": S1_additionalFiguresTables

**SUPPLEMENTARY MATERIAL 1. Complementary figures and tables from: DISPERSAL AND PLANT ARRANGEMENT CONDITION THE TIMING AND MAGNITUDE OF COFFEE RUST INFECTION**

Emilio Mora Van Cauwelaert, Cecilia González González, Denis Boyer, Zachary Hajian Forooshani, John Vandermeer , Mariana Benítez

### 1. Equilibrium points

The equilibrium points of the model with $m$= 0 (no diffusion) are:

$$(S_{eq}, I_{eq}, X_{eq}) \in \{(1,0,0), (\frac{1}{R_0}, \frac{\rho}{\gamma}(1-\frac{1}{R_0}), \frac{\alpha}{\mu} I_{eq})\}$$

For $R_0$ defined as:

$$R_0 = \frac{\beta_2 + \frac{\beta_1 \alpha}{\mu}}{\gamma}$$

The value of $R_0$ determines the stability for both equilibria. When $R_0$ is less than 1, the first equilibrium (1,0,0) is stable and the second equilibrium ($1/R_0$, $\rho/\gamma$ ($1-1/R_0$), $\alpha/\mu$ $I_{eq}$) is unstable. After any initial perturbation, the system reaches the first equilibrium where there is no infection. When $R_0 \geq 1$, the first equilibrium point becomes unstable and the system reaches the second equilibrium (now stable) point where $S$ decreases with $R_0$, whereas $I$ and $X$ "invade", or increase with $R_0$ (see (Cunniffe and Gilligan, 2010) for the full linear analysis).

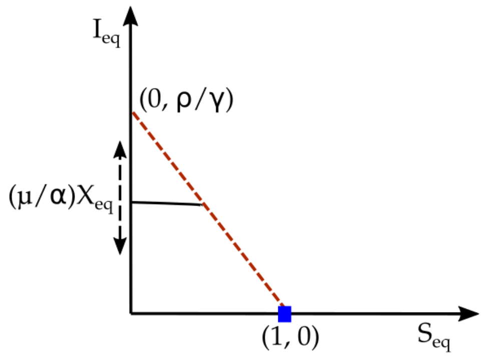


**Fig. S1.1 Relationship between the equilibrium points of the model ($S_{eq}$, $I_{eq}$, $X_{eq}$) and the parameters.** We represent $I_{eq}$ in function of $S_{eq}$ and represent the value of $X_{eq}$ as a slider in the $I_{eq}$ axis. The blue dot represents the first equilibrium point (stable when $R_0 < 1$) and the red dotted line represents the possible values of $S_{eq}$ and $I_{eq}$ in the second equilibrium point (stable when $R_0 \geq 1$).

### 2. *S*, *I* and *X* dynamics

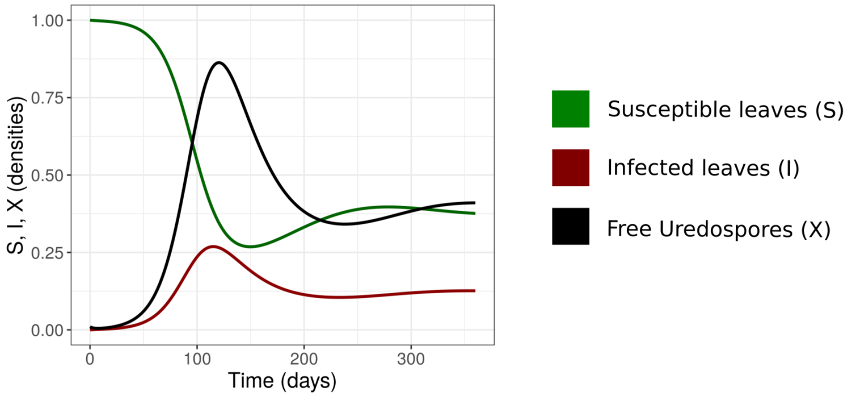


**Fig. S1.2 Example of infection dynamics for susceptible, infected leaves and free uredospores.** For this simulation, $\alpha$=0.65, $\beta_1$= $\beta_2$= 0.035, $\rho$= 0.011, $\mu$= 0.2, $\gamma$=0.056 and $I_0$ = 0.01

### 3. MATI difference with $I_{eq}$ for the single plant

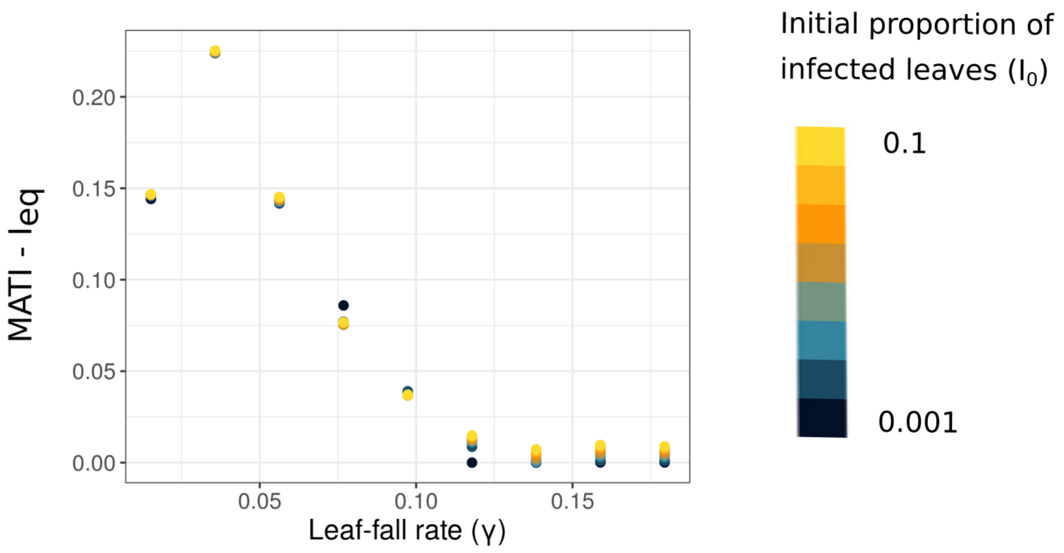


**Fig. S1.3. Difference between MATI and I at equilibrium (Ieq) in isolated coffee plants as a function of the leaf-fall rate ($\gamma$), with different initial proportion of infected leaves ($I_0$).**

The difference between MATI and the final amount of infected leaves in equilibrium ($I_{eq}$) shows the same dependence on $\gamma$ (Fig. S1.3; see Fig. S1.1 for the influence of $\gamma$ on equilibria and stability).

### 4. MATI dependence on distance-between-plant index <H>

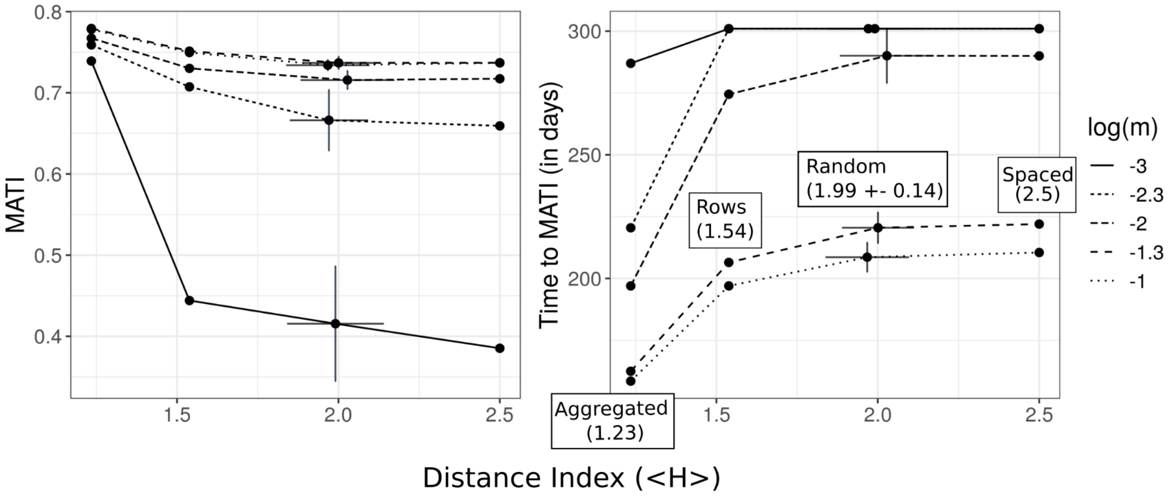


**Fig. S1.4. MATI and time to MATI as a function of the distance-between-plant index <H>.** Values of MATI and time to MATI corresponding to four values of <H>, with 2 levels of diffusion (high: 0.01 and low: 0.001) and one level of $I_0$ (0.001) (the figure for $I_0$ = 0.1 is very similar). For the random arrangement, each point represents a 30-simulation average, and the error bars the standard deviation both in the <H> and in MATI and time to MATI. The values of $\alpha$, $\beta_1$, $\beta_2$, $\rho$ and $\mu$ are shown in Table 1. We used $\gamma$=0.015.

### 5. Individual tree heterogeneity

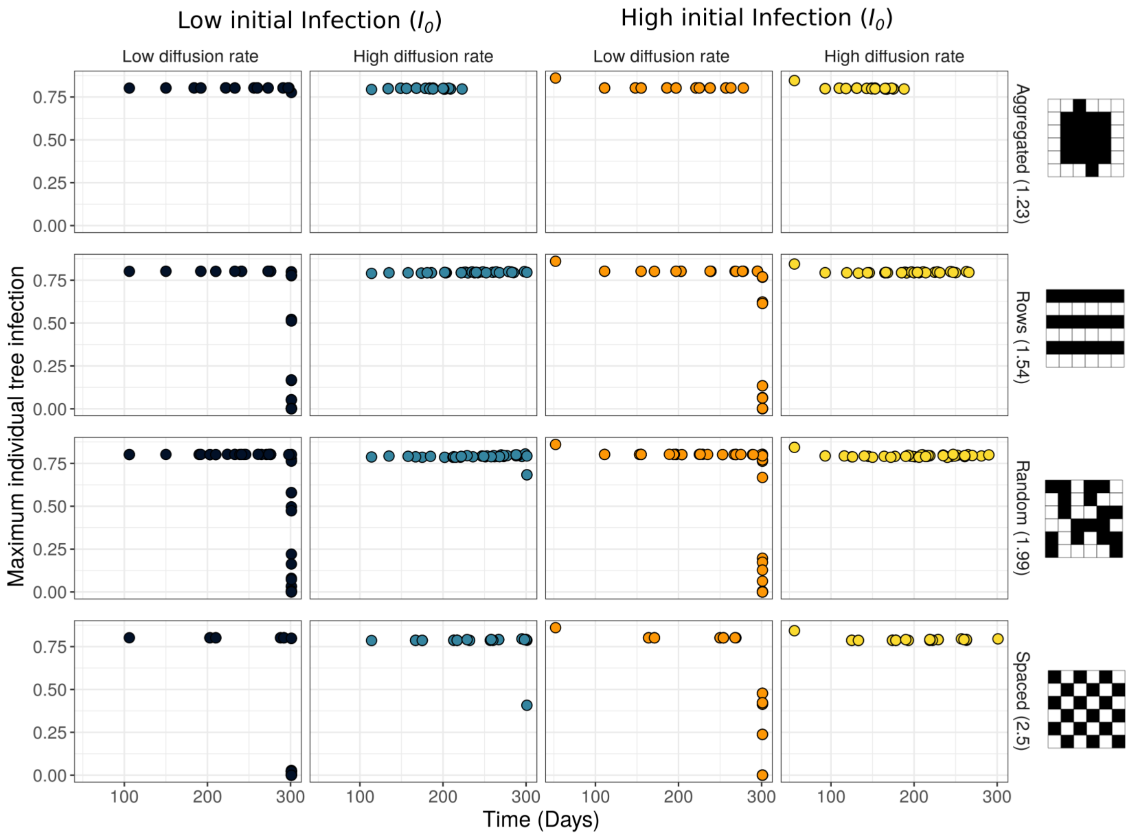


**Fig. S1.5 Maximum individual tree infections and the time to reach it.** This figure represents each tree's values for the different plant arrangements (aggregated, random, rows and spaced), two levels of diffusion (low=0.001 , high= 0.01) and two levels of $I_0$ (low= 0.001, high= 0.1). The values of $\alpha$, $\beta_1$, $\beta_2$, $\rho$ and $\mu$ are shown in Table 1. We used $\gamma$=0.015.

### 6. Time lags measurements from real time series and from simulations.

**Table. S1. 1** Duration of the time lag (Time Lag, TL), the growth phase per se (Time Growth, TG) and the time to MATI (TL+ TG) measured in 5 different plots subject to coffee rust infections (Fig. 2). PP: Precipitation

| Plot Letter in Fig.2 | Day when PP> 50mm | Time Lag (TL) | Time Growth (TG) | Time to MATI (TL + TG) |
|---|---|---|---|---|
| B | 67 | 24.5 | 104.5 | 129 |
| C | 114 | 80 | 105 | 185 |
| C | 465 | 145.5 | 180.5 | 326 |
| F | 105 | 81.5 | 120.5 | 202 |
| G | 104 | 134.5 | 132.5 | 267 |
| I | 896 | 27 | 68 | 95 |

**Table. S1. 2** Duration of the time lag (Time Lag, TL), the growth phase per se (Time Growth, TG) and the time to MATI (TL+ TG) from simulations, for the different combinations of planting arrangements, diffusion rate (*m*) and initial proportion of infected leaves ($I_0$). For the random arrangement the values are the mean and standard deviation (+-) of 30 replicates.

| Arrangement | m | *I0* | Time Lag | Time Growth | Time to MATI (TL + TG) |
|---|---|---|---|---|---|
| Aggregated (1.23) | 0.001 | 0.001 | 99 | 188 | 287 |
| | | 0.1 | 61 | 191 | 252 |
| | 0.005 | 0.001 | 78 | 143 | 220 |
| | | 0.1 | 42 | 144 | 186 |
| | 0.01 | 0.001 | 73 | 124 | 197 |
| | | 0.1 | 33 | 130 | 162 |
| | 0.05 | 0.001 | 69 | 94 | 162 |
| | | 0.1 | 23 | 101 | 124 |
| | 0.1 | 0.001 | 69 | 89 | 158 |
| | | 0.1 | 22 | 94 | 116 |
| Rows (1.54) | 0.001 | 0.001 | 150 | 150 | 300 |
| | | 0.1 | 113 | 187 | 300 |
| | 0.005 | 0.001 | 94 | 206 | 300 |
| | | 0.1 | 60 | 226 | 285 |
| | 0.01 | 0.001 | 84 | 190 | 274 |
| | | 0.1 | 50 | 189 | 239 |
| | 0.05 | 0.001 | 83 | 123 | 206 |
| | | 0.1 | 34 | 132 | 166 |
| | 0.1 | 0.001 | 88 | 109 | 197 |
| | | 0.1 | 33 | 118 | 150 |
| Random (1.99) | 0.001 | 0.001 | 135 (+- 38) | 166 (+- 38) | 300 (+- 0) |
| | | 0.1 | 96 (+- 30) | 205 (+- 30) | 300 (+- 0) |
| | 0.005 | 0.001 | 102 (+- 19) | 199 (+- 19) | 300 (+- 0) |
| | | 0.1 | 70 (+-20) | 224 (+- 21) | 294 (+- 10) |
| | 0.01 | 0.001 | 86 (+- 15) | 203 (+- 19) | 290 (+- 11) |
| | | 0.1 | 56 (+- 17) | 201 (+- 18) | 257 (+- 12) |
| | 0.05 | 0.001 | 83 (+- 8) | 137 (+- 8) | 221 (+- 16) |
| | | 0.1 | 34 (+- 7) | 146 (+- 8) | 179 (+- 5) |
| | 0.1 | 0.001 | 88 (+- 6) | 120 (+- 5) | 209 (+- 6) |
| | | 0.1 | 32 (+- 5) | 129 (+- 6) | 161 (+- 5) |
| Spaced (2.5) | 0.001 | 0.001 | 147 | 153 | 300 |
| | | 0.1 | 108 | 192 | 300 |
| | 0.005 | 0.001 | 118 | 182 | 300 |
| | | 0.1 | 78 | 216 | 294 |
| | 0.01 | 0.001 | 108 | 182 | 290 |
| | | 0.1 | 67 | 185 | 252 |
| | 0.05 | 0.001 | 92 | 130 | 222 |
| | | 0.1 | 44 | 135 | 179 |
| | 0.1 | 0.001 | 96 | 115 | 210 |
| | | 0.1 | 38 | 123 | 161 |

### 7. Sensitivity analysis (size of the integration step and plant-level parameters variation)

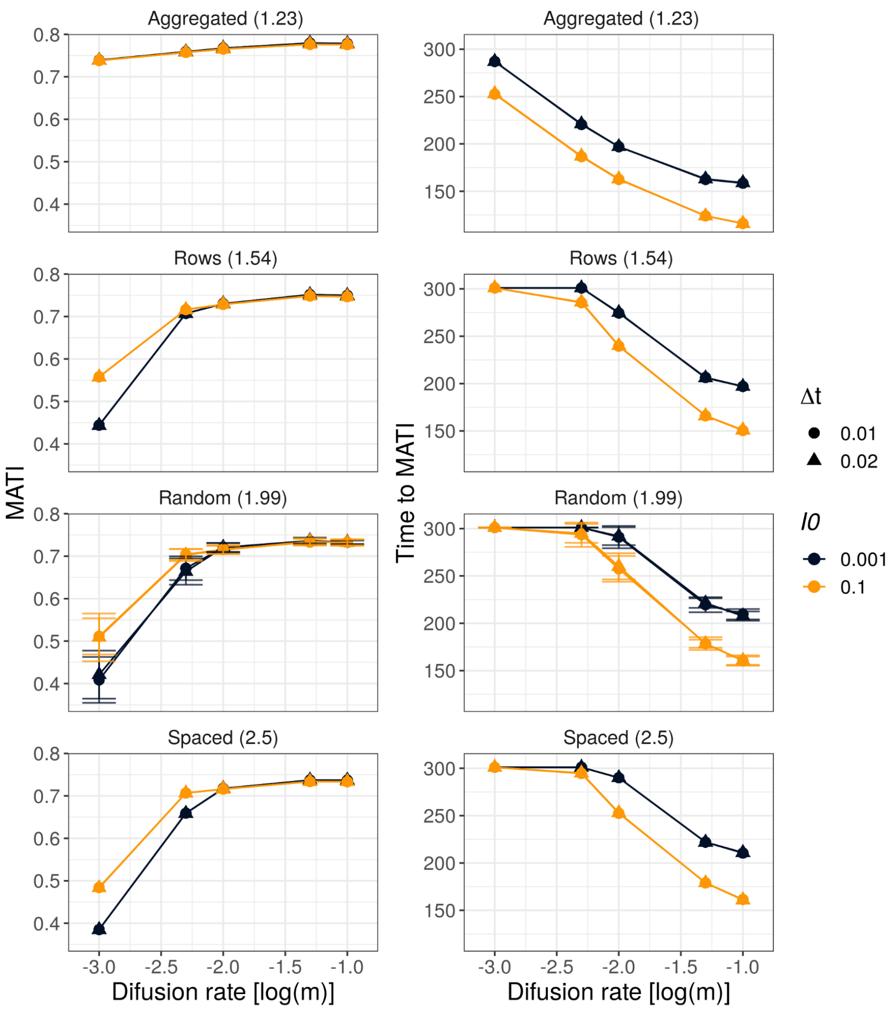


**Fig. S1.6. Robustness of MATI and time to MATI results for two values of the integration step (Δt).** Simulations of four different planting arrangements (aggregated, random, rows and spaced) with five levels of diffusion rate (*m* between 0.001 and 0.1, or log(m) between -3 and -1), two levels of $I_0$ (0.001 and 0.1, dark and orange lines respectively), and two values of Δt (0.01, 0.02). For the random arrangement, each point represents a 30-simulation average, and the error bars the standard deviation. The values of $\alpha$, $\beta_1$, $\beta_2$, $\rho$ and $\mu$ are shown in Table 1. We used $\gamma$=0.015.

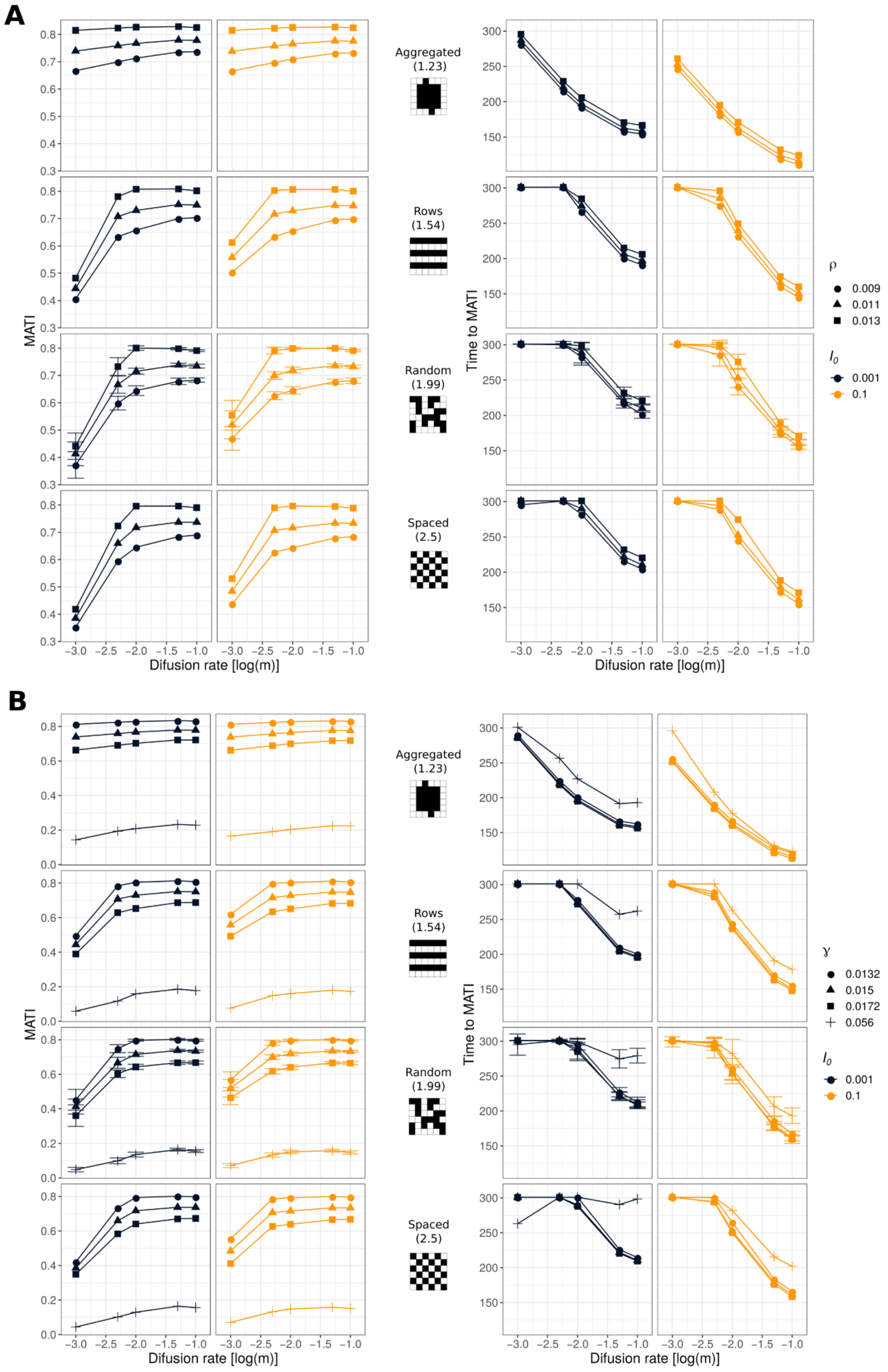

A
B
Aggregated (1.23)
Rows (1.54)
Random (1.99)
Spaced (2.5)
MATI
Time to MATI
Difusion rate [log(m)]
ρ
0.009
0.011
0.013
γ
0.0132
0.015
0.0172
0.056
$I_0$
0.001
0.1

C

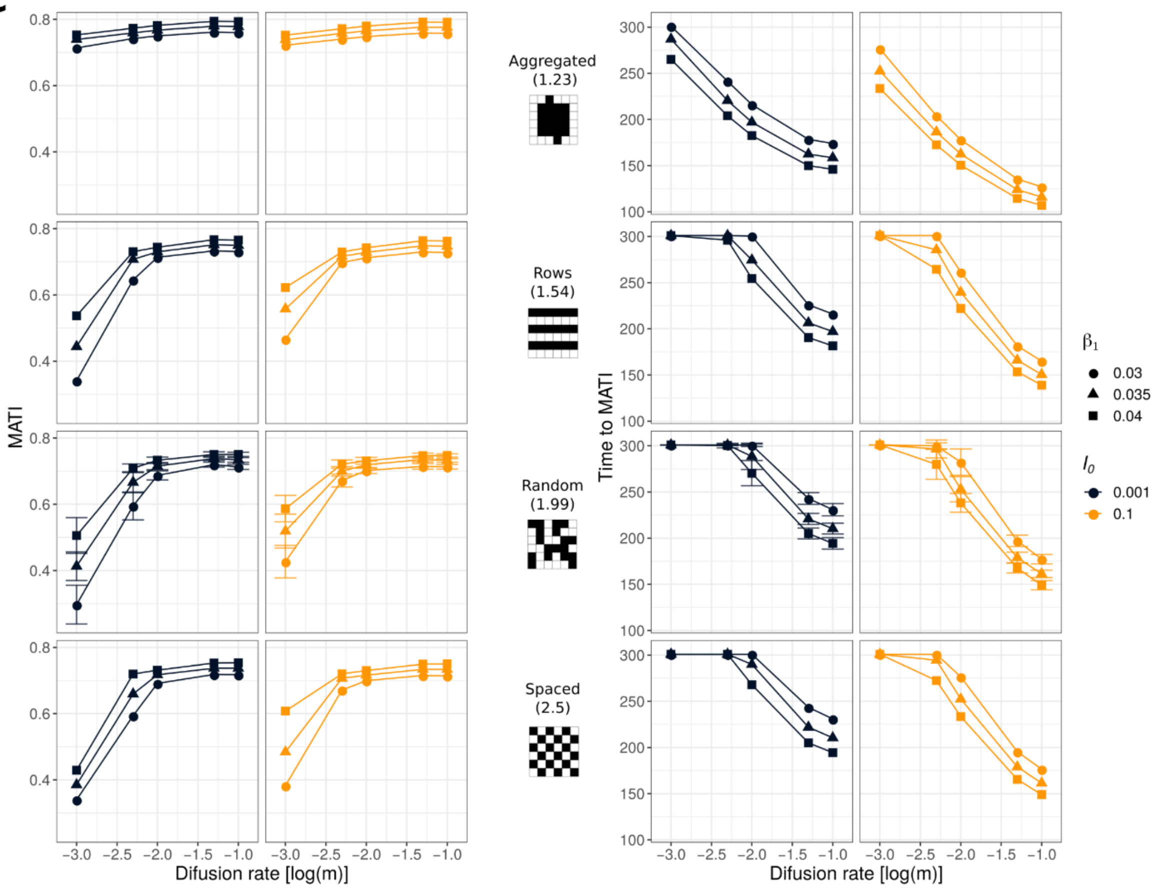

MATI
Aggregated (1.23)
Rows (1.54)
Random (1.99)
Spaced (2.5)
Time to MATI
β1
0.03
0.035
0.04
I0
0.001
0.1
Difusion rate [log(m)]


D

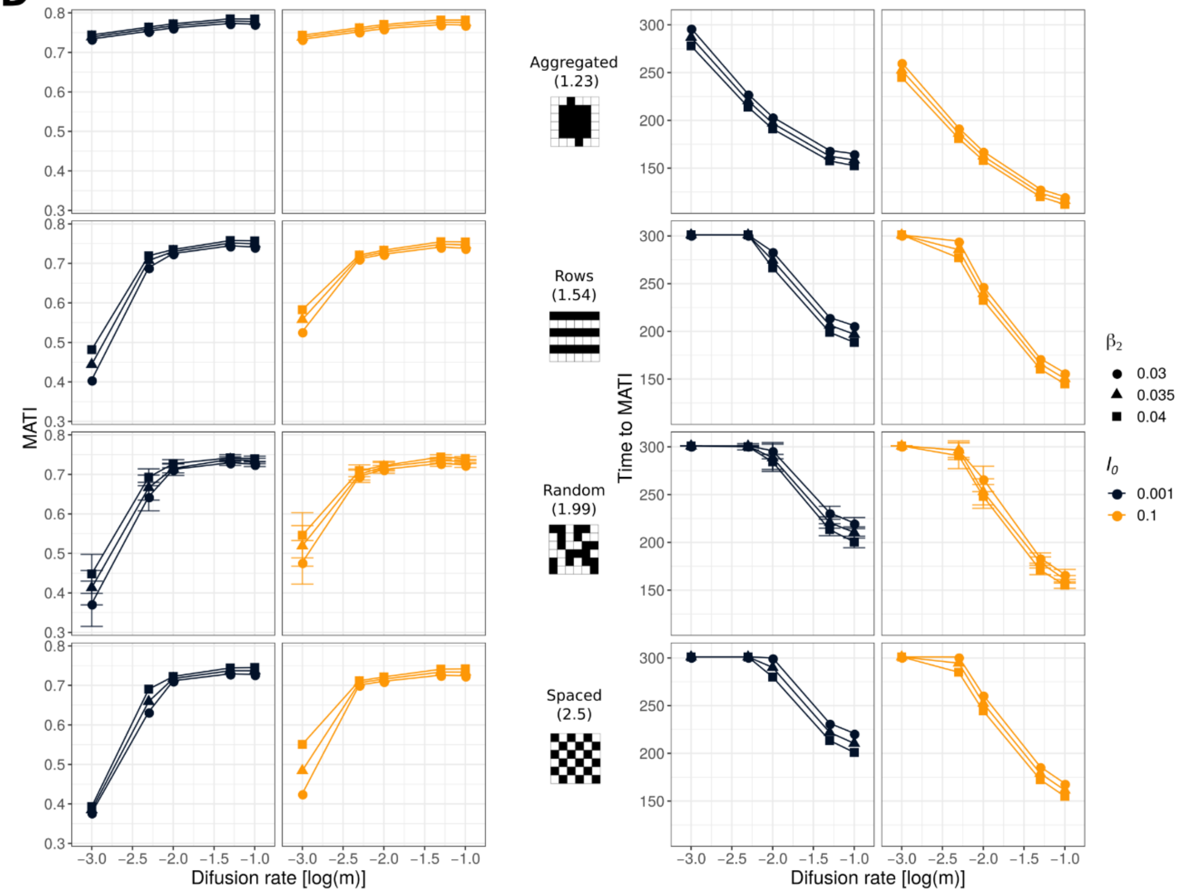

MATI
Aggregated (1.23)
Rows (1.54)
Random (1.99)
Spaced (2.5)
Time to MATI
β2
0.03
0.035
0.04
I0
0.001
0.1
Difusion rate [log(m)]

E

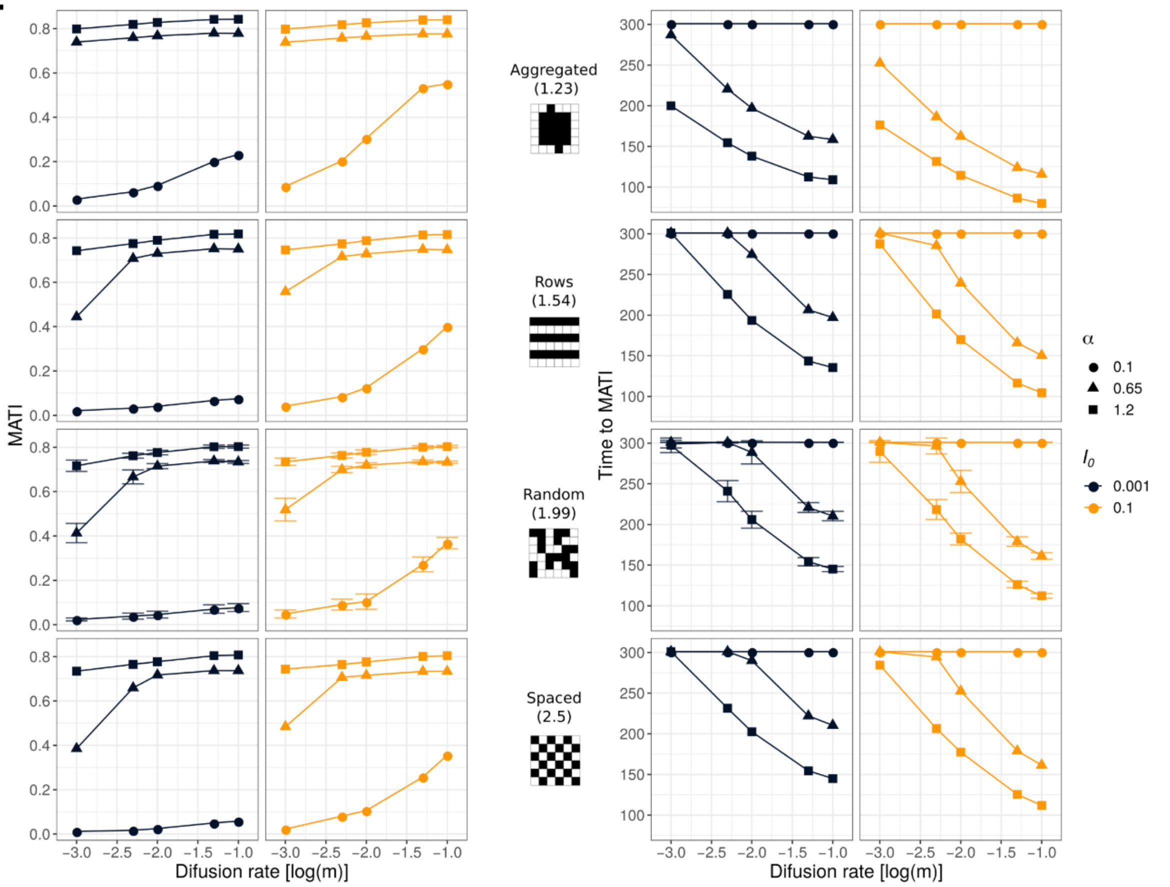
MATI
Time to MATI
Aggregated
(1.23)
Rows
(1.54)
Random
(1.99)
Spaced
(2.5)
α
0.1
0.65
1.2
$I_0$
0.001
0.1
Difusion rate [log(m)]
Difusion rate [log(m)]

F

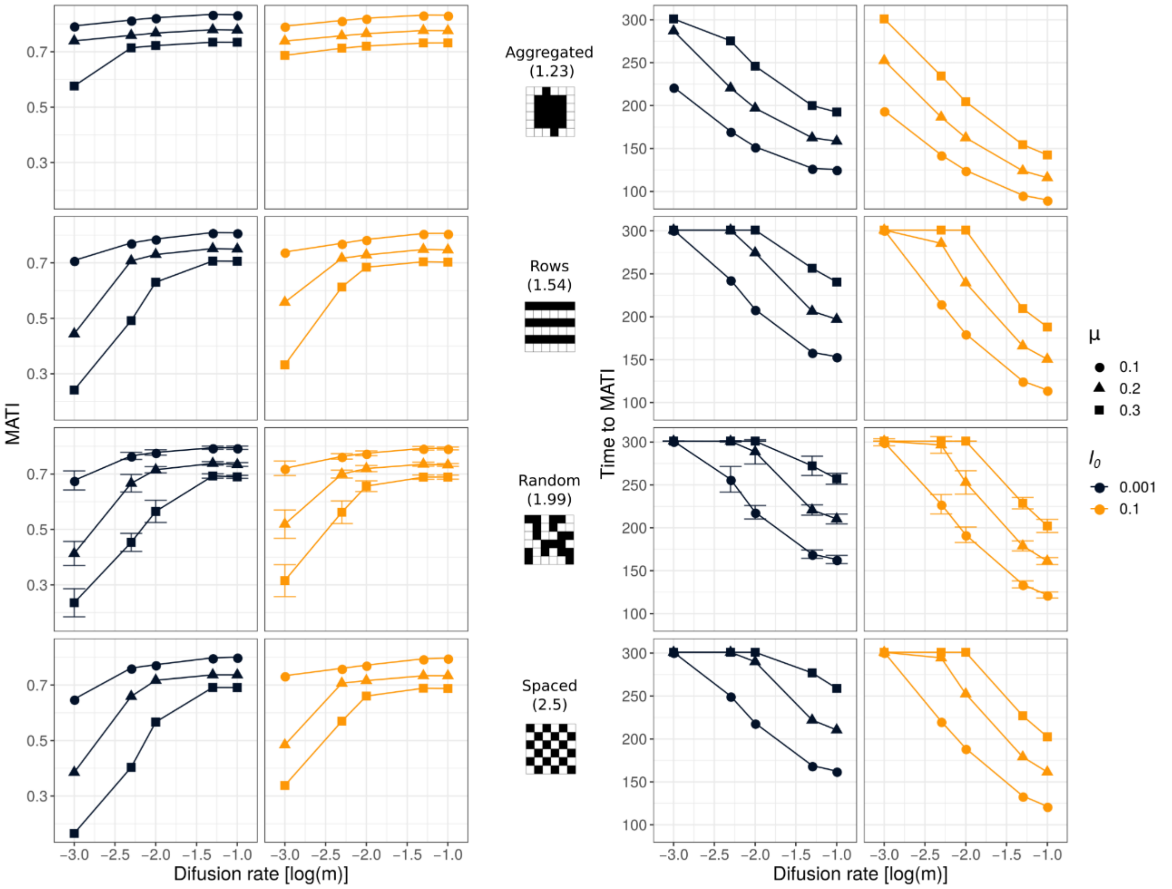
MATI
Time to MATI
Aggregated
(1.23)
Rows
(1.54)
Random
(1.99)
Spaced
(2.5)
μ
0.1
0.2
0.3
$I_0$
0.001
0.1
Difusion rate [log(m)]
Difusion rate [log(m)]

**Fig. S1.7. Robustness of MATI and time to MATI results for different values of the plant-level parameters.** We represent the variation in the results of Fig. 4 (main manuscript) produced by changing one parameter at a time within its estimated range (A:$\rho$, B: $\gamma$ , C: $\beta_{1,}$, D: $\beta_2$, E: $\alpha$, F: $\mu$). For each scenario we represent the results with the used values of the parameters in the main manuscript (triangles) and with the lowest and highest values of the estimated range (circle and square). In the γ scenario we show the results with a higher value (γ= 0.056, cross point) that accounts for leaf-removal practices. The scenarios represent the four different planting arrangements (aggregated, random, rows and spaced) with five levels of diffusion rate (*m* between 0.001 and 0.1, or log(m) between -3 and -1) and two levels of $I_0$ (0.001 and 0.1, dark and orange lines respectively). For the random arrangement, each point represents a 30-simulation average, and the error bars the standard deviation.
