## Supplementary material for "Dispersal and Plant Arrangement Condition the Timing and Magnitude of Coffee Rust Infection": S3_parameterEstimation

### 0. The model

The diagram for the local plant model is:

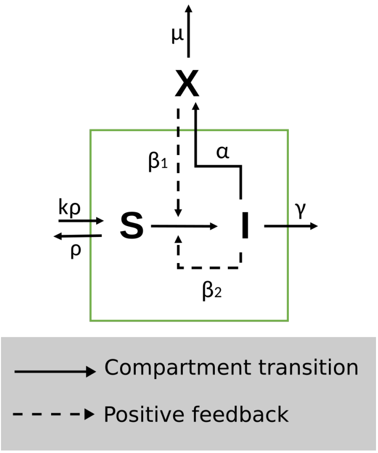


This model is then described by the following set of ODE. As we only focus on one plant, we removed the indices of the equation presented in the main text in order to simplify the notation.

$$\begin{aligned} \frac{dS}{dt} &= \rho(K - S) - \beta_1 X \frac{S}{K} - \beta_2 I \frac{S}{K} \\ \frac{dI}{dt} &= \beta_1 X \frac{S}{K} + \beta_2 I \frac{S}{K} - \gamma I \\ \frac{dX}{dt} &= \alpha I - \mu X \end{aligned} \tag{1}$$

where:

$S$: number of susceptible leaves in one coffee tree.

$I$: number of infected leaves with infective spores in one coffee tree.

$X$: amount of infective packages of spores *outside* the plant (see diagram). We call them packages since 15 to 30 spores are needed per cm2 per leaf to start an infection (Bock 1962).

$K = b/\rho$ is the carrying capacity (number of leaves per plant) where $\rho$ is the inverse of the average time needed for a newly mature susceptible leaf to fall, and $b$ is the number of leaves produced per unit of time (here [t]= days). We can also express $b$ as $K\rho$ (see general diagram). Here we assume for simplicity that K = 1, so that $\rho$ is equal to $b$. $\beta_1$ and $\beta_2$ are the rates of primary and secondary infection. $1/\beta_1$ is the time

Table 1: Estimated parameters range. The letters indicate the references. a-Rakocevic and Takeshi (2018), b- Mulinge and Griffiths, E., (1974), c- Firman and Wallis (1965) d- Leguizamón-Caycedo et al. (1998), e-Bock (1962), f-Rayner (1961), g-Gagliardi et al. (2020), h- Boudrot et al. (2016), i- Silva-Acuña et al. (1999) j- Deepak, K., et al. (2012), k- Nutman, et al. (1963).

| Parameters | $\rho$ | $\gamma = \rho + h_I$ | $\beta_1 \beta_2$ | $\alpha$ | $\mu$ |
|---|---|---|---|---|---|
| Definition | Natural leaf growth and death rate | Infected leaves death rate | 1st and 2d infection rate | Recruitment rate | Spore death rate |
| Units | t-1 | t-1 | nspore-1 t-1, nleaf-1. t-1 | nspore nleaf-1. t-1 | t-1 |
| Range | [0.009-0.013] | pI= [0.0042] | [0.03-0.04] | [0.1-1.2] | [0.1-0.3] |
| Section | I, III | III | I | II | II |
| References | a, b | a, c | d, e | e, f, g, h, i | j, k |

taken for one susceptible leaf in contact with one external package of infective spores to become an infected leaf with new infective spores. In this sense, this term includes spores that are surrounding the tree. $1/\beta_2$ is the time taken for one susceptible leaf, in contact with another infective leaf, to become an infected leaf with new infective spores[1]. Since one infected leaf has multiple spores, one leaf is sufficient to start an infection into another leaf. Besides, we are assuming that dispersion of spores from one leaf to another inside the tree is included in the $\beta_2$ term. $\alpha$ is the spore recruitment from infected leaves to the outside of the plant. $\gamma$ is the infected leaf fall rate. This rate considers natural leaf fall and leaf fall due to the rust infection ($\gamma = \rho + h_I$). $1/\gamma$ is then the average time taken for an infected leaf to fall. Finally $\mu$ is the spore death rate.

**Assumptions**

The model is set up for the rainy season. This means that:

a) The spores are washed out from the top to the bottom of the plant (Bock 1962). So, spores from one leaf can be in contact with any other leaf in the plant. As the vector inside the plant is well mixed, we can assume that leaves are also "well mixed" (since $\beta_2$ includes this kind of infection).

b) Optimal conditions for rust development are always met.

### I. In field measurements of parameters (direct estimation)

#### a) $\rho$ measurement

In Rakocevic and Takeshi Matsunaga (2018) leaf life span and leaf expansion time were recorded in growing degree days (GDD). GDDs were calculated as $GDD = T_{mean} - T_b$, where $T_{mean}$ is the mean air temperature calculated as the average of daily minimum and maximum air temperatures, and $T_b$ is the base temperature for each species (Rakocevic and Takeshi Matsunaga 2018). The calculated $T_b$ is 10.2 °C. Using Rakocevic supplementary material (2018) we estimated the relationship between GDD and a normal day in active season (The active season goes from oct-13 to abr-13 and from oct-14 to dic-14) (the warm-rainy season) and reduced season (remaining days, dry-cold season). We obtained the mean of GDD/day in active and reduced season in order to tranform the reported GDD for leaf growth parameters to days.

For Active season GDD/days = 13.7, for Reduced season, GDD/days = 9.25

In rust infection, fully expanded leaves are the most infected[2]. So we consider the adult life span of a leaf (S) from the moment when it is fully expanded to the moment when it falls. With this in mind, we

[1]Assuming $K = 1$, if this is not the case, $\beta_n = \beta_n^* K$ ($n \in 1, 2$), where $1/\beta_n^*$ is the characteristic time

[2]With natural infection, young lesions have been observed on leaves of all ages except those still of juvenile (glossy) appearance. This appearance is lost on average in week 12.2 [85.4 days] of age (Rayner 1961). As the incubation period averages 5 weeks it is evident that leaves are rarely infected until fully expanded (Rayner 1961)

Table 2: Average GDD in active o reduced season. CFA refers to the climate classification, (see Rakocevic and Takeshi (2018) for full explanation)

| Bar | Type | GDD.LifeSpan | GDD.LeafExpansion | GDD.AdultLife |
|---|---|---|---|---|
| Bar1 | Act_Growth_CFA | 1809.89 | 671.03 | 1138.85 |
| Bar3 | Red_Growth_CFA | 1769.96 | 632.41 | 1137.55 |
| Bar5 | Act_Growth_CFA | 2076.05 | 806.21 | 1269.84 |
| Bar7 | Red_Growth_CFA | 1969.58 | 704.83 | 1264.75 |
| Bar9 | Act_Growth_CFA | 1876.43 | 864.14 | 1012.29 |
| Bar11 | Red_Growth_CFA | 2009.51 | 811.03 | 1198.47 |
| Bar13 | Act_Growth_CFA | 2408.75 | 840.00 | 1568.75 |
| Bar15 | Red_Growth_CFA | 2169.20 | 801.38 | 1367.82 |

Table 3: Mean life (F) range in GDD and real days

| | Act_Growth_CFA | Red_Growth_CFA |
|---|---|---|
| meanGDD_Adult.Life | 1247.43 | 1242.15 |
| sdGDD_Adult.Life | 238.63 | 98.58 |
| MinDay_Adult.Life | 73.64 | 123.63 |
| MaxDay_Adult.Life | 108.47 | 144.94 |

transformed the reported data for full life span (F), leaf expansion (E) and adult life span ($F - E$). This adult life time is then assumed to be $1/\rho$ (this measurement is more precise than considering expansion, because expansion depends a lot on the final size of the leaf). Our model simulates infection in climatic conditions with optimum precipitation and temperature values, so we only took the values for the "Active season" (warm and rainy). Specifically we only took Rakocevic's reported data for CFA Koppen-climate. This is a good proxy since during "Active Season" this climate is similar to the one present in Mexican coffee plantations in the rainy season (table 2 and 3). Specifically we took the {mean-sd, mean+sd} for GDD_Adult.Life (MinDay_Adult.Life and MaxDay_Adult.Life) and for each value, we transformed it to normal days (table 3).

In active season, the time taken for a fully expanded leaf $S$ to fall is in $[74 - 108]$ days. So, $\rho \in [0.009, 0.013]$. (For reduced season (dry season) $\rho \in [0.007 - 0.008]$.)

#### b) Primary and secondary infection measurement ($\beta_1, \beta_2$)

The average time taken for a susceptible leaf in contact with an infective leaf or a infective package of spores, to become an infected leaf with new infective spores ranges from 26 to 35 days in average in conditions similar to the ones present in mexican coffee agroecosystems (Leguizamón-Caycedo, Orozco-Gallego, and Gómez-Gómez 1998). So in this case: $\beta_1, \beta_2 \in [0.03, 0.04]$.

### II. Data-based estimation of parameters

#### a) Mortality rate of spores ($\mu$)

If there is no new infection, we can say that the amount of infective spores in time follows the equation: $X(t) = X(0)e^{-\mu t}$ in our model. Deepak et al. (2012) reported that for rust RI variety only the 4% of original spores is viable after 15 days. In each day trial, spores were germinated for 16 hours. For rust RVIII, this percentage increases to 23% (Deepak, Hanumantha, and Sreenath 2012). We can use this data to estimate the value of $\mu$ in our equation.

$$X(15) = 0.04X(0) = X(0)e^{-\mu(15)} \tag{2}$$

$$e^{-\mu(15)} = 0.04 \tag{3}$$

$$\mu = \frac{ln(0.04)}{-15} \tag{4}$$

$$\mu = 0.2 \tag{5}$$

And for RVIII

$$\mu = \frac{ln(0.23)}{-15} \tag{6}$$

$$\mu = 0.1 \tag{7}$$

In this case $\mu \in [0.1, 0.2]$

In another work by Nutman et al, (1963) spore viability is calculated at 22°C. They decrease in viability follows a power law (Nutman, Roberts, and Clarke 1963):

$$X(t) = X(0)(t+1)^{-0.809} \tag{8}$$

We generated a database with Nutman equation and fitted it to our exponential model for the first 5 days.

The $\mu$ that fitted the plot best is equal to 0.343 and is statistcally significant. We can say that $\mu \in [0.1, 0.3]$.[3] After 75 days, the viability remains at 20%. In this case $\mu = 0.02$. For models than consider spores travelling at high altitudes this might be relevant.

#### b) Recruitment rate of new spores ($\alpha$)

$\alpha$ is the rate of recruitment of a new package of infective spores (to the external surrounding medium) from one infective leaf. If we only look at the production of new spores we have the equation:

$$\frac{dX}{dt} = \alpha I(t) \tag{9}$$

In this case, $[\alpha]$ = [spores/infected leaf/day]. We know from Rayner (1961)that one lesion produces around 300 thousand spores in a period of 3 to 5 months (Rayner 1961). This means that per day, one lesion produces around 2000 to 3000 spores (assuming constant production).

One rusted leaf has from 2 to 6 active lesions (amount of lesions that are constantly producing spores). (Firman and Wallis 1965)(Silva-Acuña et al. 1999). So, one infective leaf produces from 4000 to 18 000 spores per day.

[3] In both estimations, temperature is constant, but we know that this death rate decreases a lot with very low temperatures (4°C) (Capucho et al. 2005)

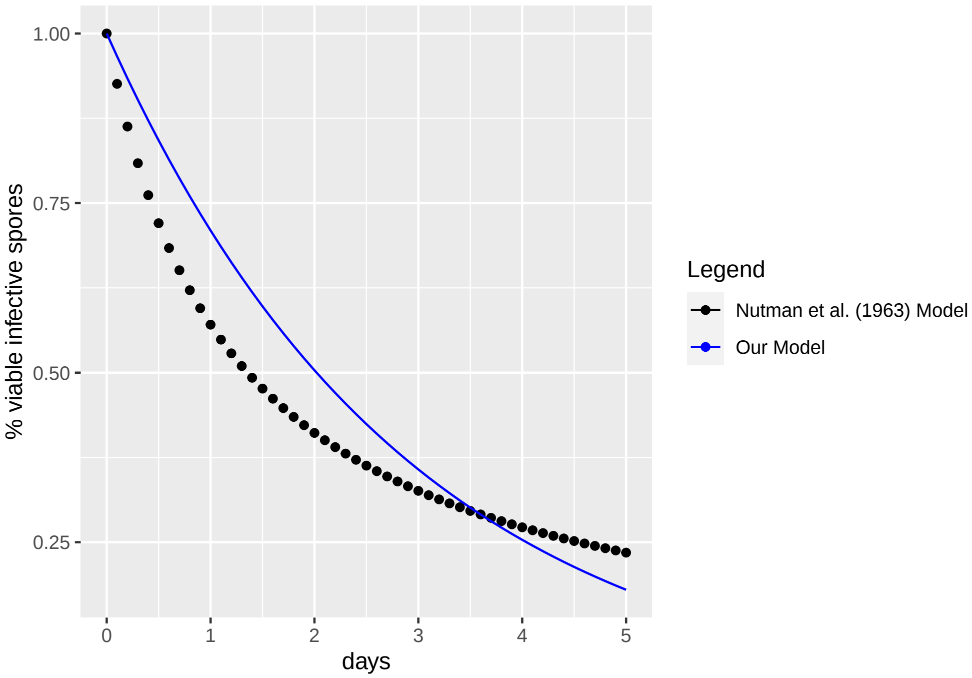


Figure 1: Viability of uredospores (Nutman, 1963) and our model fitting

According to Boudrot et al. (2016) and Gagliardi et al. (2020) there is a 1000-1 relationship between spores in the leaf and spores in the air (Boudrot et al. 2016) (Gagliardi et al. 2020) . As $X$ represent specifically spores in the air surrounding the tree (that eventually fall down to the tree) we will use this relation.

Finally we know that, in order to be infective, the optimum quantity of spores (or package of spores) is between 15 to 30 spores (Bock, 1962). Taking all these factors into account we can conclude that one infected leaf releases from 0.1 to 1.2 infective packages of spores per day to the air. We will asume this same quantity falls back into the tree.

$\alpha \in [0.1, 1.2]$ spores/rusted leaf/day

### III. Data based fitting-estimation

We used time-series data and few assumptions on the behaviour of some variables to extract the range of the missing parameters or to confirm the already reported ones. For this we extracted data using WebPlot-digitizer. We first used the data from Firman and Wallis (1965) on the fallen rusted and non rusted leaves to estimate $\rho$ and $\gamma$ (Firman and Wallis 1965). We then used Mulinge and Griffiths (1974) data set to have another estimation of $\rho$ (Mulinge and Griffiths 1974).

**a) Relationship between $\rho$ and $\gamma$**

With data from Firman and Wallis (1965) we first built a table (Table 4) with the total number of susceptible and infected leaves in a tree per month (roughly) from october 1961 to july 1963. The method was the following:

The paper reports the total number of fallen leaves per month (F) (Fig3. in the original paper) and the total amount of leaves in the tree both in october 61 and july 1963 (Table 8 in the original paper). The change in foliation during a specific interval ($\Delta$) can be described with the following equation:

$$T_{(t+\Delta)} = T_{(t)} + [P_{(t+\Delta)} - P_{(t)}] - [F_{(t+\Delta)} - F_{(t)}] \tag{10}$$

where P is the new produced leaves, F, the fallen leaves and T, the total number of leaves in the tree. So, we can write:

$$[P_{(t+\Delta)} - P_{(t)}] = T_{(t+\Delta)} - T_{(t)} + [F_{(t+\Delta)} - F_{(t)}] \tag{11}$$

Firman and Wallis (1965) report that $T_{(jul63)} = 3613$, $T_{(oct61)} = 4000$ and that the total amount of fallen leaves during that period equals: $F_{(jul63)} - F_{(oct61)} = 7287$. This means that, according to equation 11, the production of leaves from october 61 to november 63 ($P_{(nov63)} - P_{(oct61)}$) equals 6900.

Now, if we assume that leaf production is constant during that time[4] (this time is 574 days) we can derive that the production of leaves per day equals: $6900/574 = 12.02$.

With this, we estimated the production of new leaves per month ($P_{t+d} - P_t$) and the total number of leaves in the tree ($T_t$) per month (Table 4). Finally, we calculated, among those total leaves ($T_t$), the total quantity of susceptible (St) and infected leaves (It) per recorded time using Fig 1 values in (Firman and Wallis 1965)] that report the proportion of infected leaves (Prop(It)) per month (Table 4).

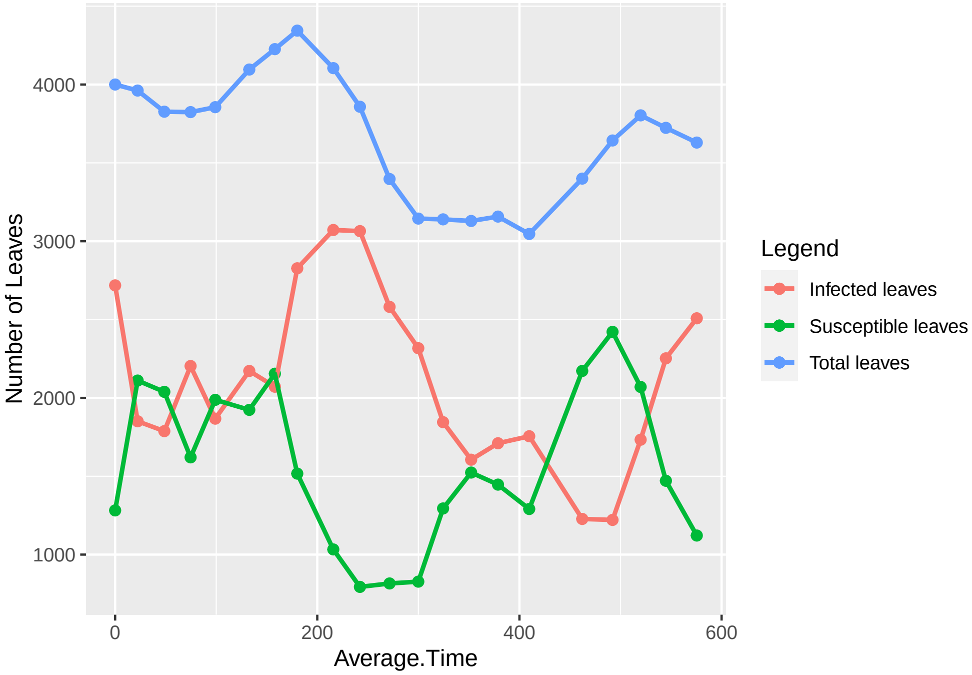


Figure 2: Proportion of Infected, Susceptible and Total Leaves. Data from: (Firman and Wallis, 1965)

[4]This makes sense since, with no infection, $\frac{dS}{dt} = b - hS = h(\frac{b}{h} - S) = \rho(K - S)$ which is the monomolecular model

Table 4: Proportion of rusted and non-rusted fallen leaves per time. Ft+d-Ft: Difference in fallen leaves, Ft: total fallen leaves, Prop(It): proportion of tree-level rust infection, Pt+d-Pt: produced leaves per recorded time, Tt: total leaves in the tree, It: number of infected leaves, St= number of susceptible leaves

| Date | Ft+d-Ft | Ft | Prop(It) | Average Time | Pt+d-Pt | Tt | It | St |
|---|---|---|---|---|---|---|---|---|
| Oct61 | NA | NA | 67.95 | 0.00 | NA | 4000.00 | 2718.03 | 1281.97 |
| Nov61 | 305.94 | 305.94 | 46.72 | 22.25 | 267.44 | 3961.50 | 1850.86 | 2110.63 |
| Dec61 | 452.32 | 758.26 | 46.72 | 48.66 | 317.43 | 3826.61 | 1787.84 | 2038.77 |
| Ene62 | 314.89 | 1073.16 | 57.62 | 74.63 | 312.23 | 3823.95 | 2203.47 | 1620.48 |
| Feb62 | 262.22 | 1335.38 | 48.44 | 99.06 | 293.61 | 3855.34 | 1867.63 | 1987.71 |
| Mar62 | 164.27 | 1499.65 | 53.03 | 132.72 | 404.57 | 4095.64 | 2172.03 | 1923.61 |
| Apr62 | 173.76 | 1673.41 | 49.02 | 158.01 | 303.93 | 4225.81 | 2071.34 | 2154.47 |
| May62 | 148.57 | 1821.98 | 65.08 | 180.19 | 266.63 | 4343.87 | 2827.08 | 1516.80 |
| Jun62 | 668.01 | 2489.99 | 74.84 | 215.85 | 428.68 | 4104.55 | 3071.68 | 1032.87 |
| Jul62 | 562.89 | 3052.88 | 79.43 | 242.18 | 316.51 | 3858.17 | 3064.40 | 793.77 |
| Aug62 | 813.71 | 3866.60 | 75.98 | 271.50 | 352.32 | 3396.78 | 2580.99 | 815.78 |
| Sep62 | 594.11 | 4460.71 | 73.69 | 299.93 | 341.82 | 3144.48 | 2317.12 | 827.36 |
| Oct62 | 299.81 | 4760.52 | 58.77 | 324.45 | 294.75 | 3139.42 | 1845.05 | 1294.37 |
| Nov62 | 344.10 | 5104.63 | 51.31 | 352.23 | 333.85 | 3129.17 | 1605.62 | 1523.55 |
| Dec62 | 291.30 | 5395.92 | 54.18 | 378.79 | 319.28 | 3157.15 | 1710.56 | 1446.60 |
| Ene63 | 482.26 | 5878.18 | 57.62 | 409.69 | 371.40 | 3046.29 | 1755.36 | 1290.93 |
| Mar63 | 277.08 | 6155.26 | 36.11 | 462.13 | 630.33 | 3399.55 | 1227.46 | 2172.09 |
| Apr63 | 117.06 | 6272.32 | 33.52 | 492.11 | 360.38 | 3642.86 | 1221.26 | 2421.61 |
| May63 | 173.79 | 6446.11 | 45.57 | 519.91 | 334.08 | 3803.15 | 1733.24 | 2069.91 |
| Jun63 | 379.93 | 6826.05 | 60.49 | 544.88 | 300.23 | 3723.45 | 2252.38 | 1471.07 |
| Jul63 | 461.45 | 7287.50 | 69.10 | 575.46 | 367.51 | 3629.51 | 2507.93 | 1121.58 |

Secondly, we used this table to estimate $\rho$ and $\gamma$. Firman and Wallis (1965) state that from May 1962 to September 1962, 87.6% of fallen leaves were rust-infected and the rest were without rust. This means that for the total amount of 2787 fallen leaves[5] in this period, 2441 were rust infected ($R_I$) and 346 were free of infection ($R_S$). We know that the amount of fallen susceptible leaves follows the next equation:

$\frac{dR_S}{dt} = \rho S$

This means that we can aproximate the value of $\rho$ by:

$$R_S(t) = \rho \int_0^t S$$

We estimated the Area under the Curve (AUC) of S from May 1962 to September 1962 and calculated the corresponding $\rho$. This gives $\rho = 0.00300$

Then, we used the same procedure to estimate $\gamma$, using AUC of I(t) and the value of fallen leaves in may62-sep62. According to the next equation:

$$R_i(t) = \gamma \int_0^t I$$

We have $\gamma = 0.0072$. So, for this system $\gamma = \rho + 0.0042$.

**b) Comparison of the estimated $\rho$ value with data fitting**

If we assume no infection, our model for the change of $S$ becomes:

$$\frac{dS}{dt} = \rho(K - S) \tag{12}$$

whose solution is:

$$S(t) = K - e^{-\rho t}(K - S_0). \tag{13}$$

This model of growth (monomolecular growth) depends on the value of $\rho$ (Fig.3). Interestingly, this explicit equation enables us to estimate $\rho$ value using a non infected data set of leaf number, if we know the value of $K$ and $S$ in time, or if $S/K$ is reported.

In Mulinge and Griffiths (1974), the value of $S/K$ and $I/K$ is reported (Fig.4). We do not have the amount of fallen leaves, so from this time series we consider a time interval where $I/K$ is close to 0 (we assume that $X/K$ is also near to zero). That is, between day 290 and day 370 which might correspond to the dry season.

From eq (12) one obtains:

$$S(t)/K = s(t) = 1 - (1 - s(0))e^{-\rho t} \tag{14}$$

$$ln(\frac{1 - s(t)}{1 - s(0)}) = -\rho t \tag{15}$$

and adjust a linear model to estimate $\rho$ (Fig.5). (The assumptions of homoscedasticity and normality of residuals are respected). We got $\rho = 0.008$ with a $R^2 = 0.99$. This value for $\rho$ is within the range estimated for non-rain season in section I ($\rho \in [0.007, 0.008]$). We will not use this result to estimate the range of $\rho$since our model is set for the rainy season, but it validates the direct method to estimate $\rho$ range presented in section I.

[5]It does not correpond exactly to what Firman reported (2616 total fallen leaves). But this difference arises from rounding and from assuming constant production of leaves. The difference is not significant.

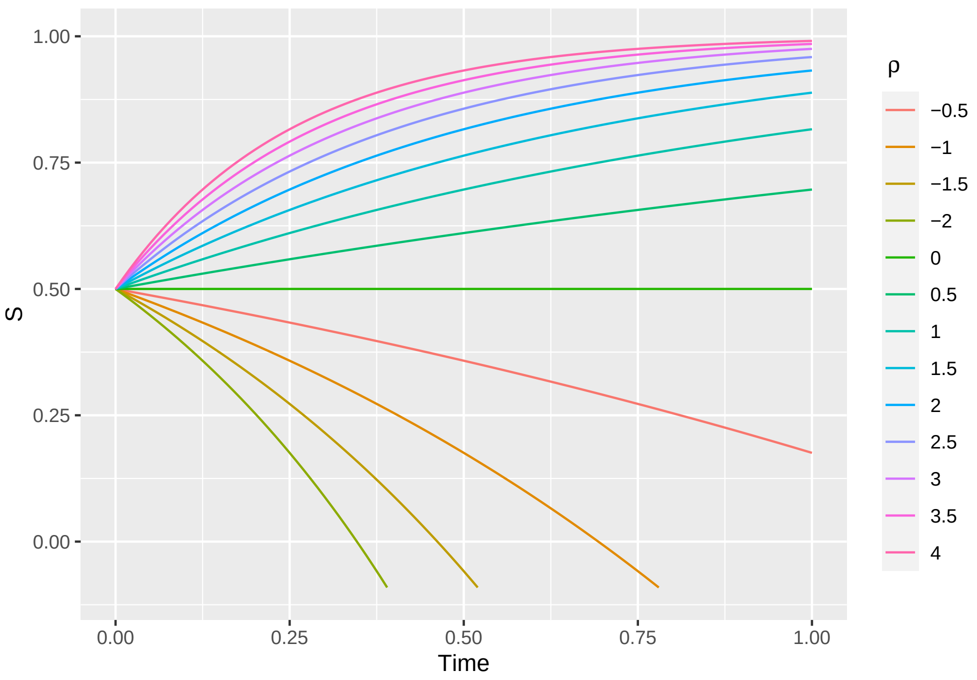


Figure 3: Time evolution of S, with $\rho$ varying from -2 to 4. $S_0$= 0.5, $K = 1$.

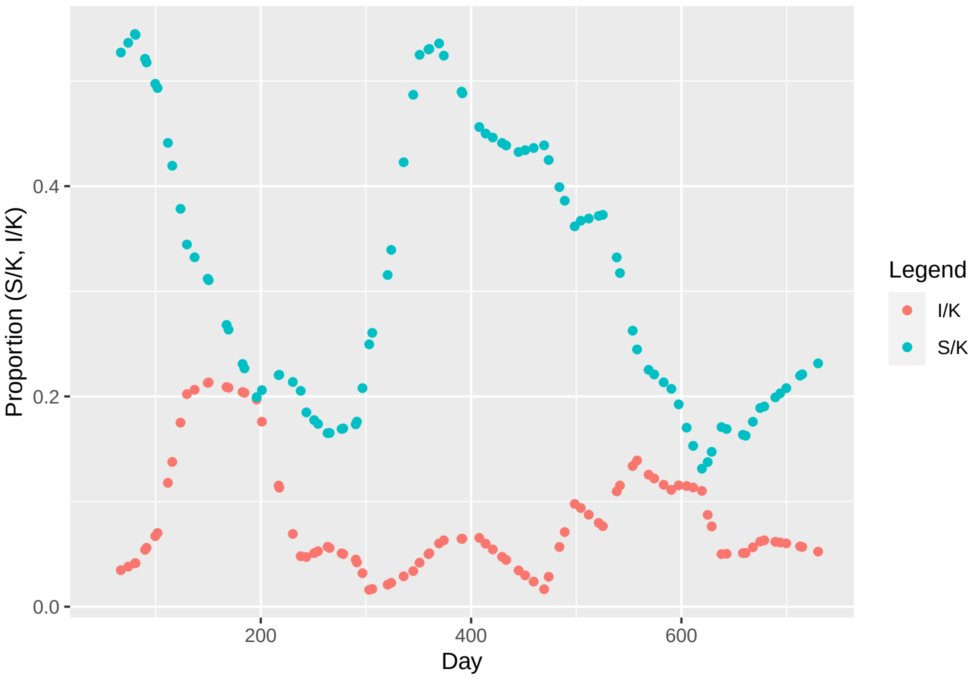


Figure 4: Proportion of infected and susceptible leaves in two epidemic years (Mulinge and Griffiths, 1974). $S$:Susceptible leaves, $I$:Infected Leaves, $K = 20$

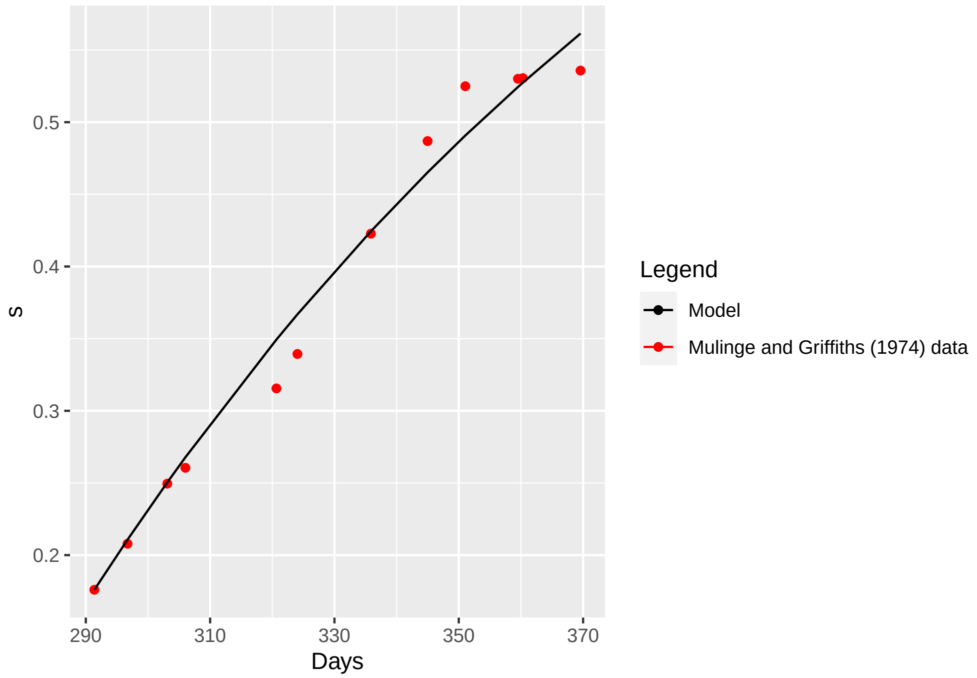


Figure 5: Linear fitting to Mulinge and Griffiths Data (1974).
